# Transcriptomic analysis of pubertal and adult virgin mouse mammary epithelial and stromal cell populations

**DOI:** 10.1101/2024.01.11.575148

**Authors:** Nika Heijmans, Katrin E. Wiese, Jos Jonkers, Renée van Amerongen

**Affiliations:** Developmental, Stem Cell and Cancer Biology, Swammerdam Institute for Life Sciences, University of Amsterdam, Science Park 904, 1098 XH Amsterdam, the Netherlands; Present address: Wageningen Bioveterinary Research, Wageningen University & Research, Lelystad, The Netherlands; Division of Molecular Pathology, Oncode Institute, Netherlands Cancer Institute, Plesmanlaan 121, 1066 CX Amsterdam, the Netherlands

**Keywords:** RNA sequencing, FACS, estrous cycle, Wnt4, estrogen, progesterone

## Abstract

Conflicting data exist as to how mammary epithelial cell proliferation changes during the reproductive cycle. To study the effect of endogenous hormone fluctuations on gene expression in the mouse mammary gland, we performed bulk RNAseq analyses of epithelial and stromal cell populations that were isolated either during puberty or at different stages of the adult virgin estrous cycle. Our data confirm prior findings that proliferative changes do not occur in every mouse in every cycle. We also show that during the estrous cycle the main gene expression changes occur in adipocytes and other stromal cells. Finally, we present a comprehensive overview of the *Wnt* gene expression landscape in different mammary gland cell types in pubertal and adult mice. This work contributes to understanding the effects of physiological hormone fluctuations and locally produced signaling molecules on gene expression changes in the mammary gland during the reproductive cycle and should be a useful resource for future studies investigating gene expression patterns in different cell types across different developmental timepoints.

## Introduction

One of the risk factors for breast cancer is an increased number of menstrual cycles, especially for every year younger at menarche [1]. The molecular basis for this remains unknown, but it likely reflects the accumulation of mutations due to a higher total number of (stem) cell divisions that underlie recurrent changes in tissue organization. The mouse provides a good model system to study the dynamics of mammary gland development and homeostasis as, similar to the human breast, it undergoes major changes in growth, differentiation and morphology during puberty and the adult reproductive cycle under the influence of ovarian hormones and locally produced paracrine signaling factors [2].

Ductal morphogenesis in puberty is mainly stimulated by estrogen [3,4]. Dynamic growth and differentiation in the adult are most obvious during pregnancy and lactation, when extensive side-branching occurs and secretory alveoli develop in response to progesterone and prolactin [5,6]. More subtle morphological changes, however, are known to occur during the estrous cycle, which repeats itself every 4 to 5 days in mice [7–12]. This short cycle is divided into four stages: proestrus, estrus, metestrus and diestrus, which correspond to the follicular (proestrus/estrus) and luteal (metestrus/diestrus) phases of the human menstrual cycle, with corresponding fluctuations in estrogen and progesterone (**Supplementary Fig. 1**) [13–15].

Alternating estrogen and progesterone signaling activity are thought to cause continuous cycles of growth and regression of lobuloalveolar outgrowths as well as sustained tertiary side-branching of epithelial ducts over time [7,8,10–16]. However, the molecular basis of the morphological changes that occur during the estrous cycle remains poorly understood. Moreover, discrepancies exist between studies investigating these proliferative changes. Some observed an increase of epithelial cells mainly during late proestrus/estrus [7,8,16]. Others reported that outgrowth of lobuloalveolar structures mainly occurs in diestrus compared to other stages [10,11,13]. More recent work showed that proliferative expansion of the epithelial ducts does not occur in every cycle in every mouse, highlighting the complexity of proliferative heterogeneity during the estrous cycle [12].

The outgrowth and regression of epithelial ducts that has been reported, is presumably the result of an indirect effect of steroid hormones on mammary stem cell (MaSC) regulation [10,17–19]. One of the candidate factors responsible for controlling mammary stem cell behavior downstream of progesterone is *Wnt4* [10,20,21], a member of the *Wnt* gene family that encodes ligands that play a pivotal role in stem cell self-renewal and tissue homeostasis in multiple different tissues [22], including the mammary gland [23–30]. Studies investigating the link between steroid hormones and *Wnt* genes or MaSC activity are often performed using ovariectomized mice receiving treatment with progesterone and/or estrogen pellets or injections [10,11,17,20,31]. Therefore, a need still exists for studies that examine the direct molecular consequences of hormonal fluctuations in the mammary gland under physiological conditions. Furthermore, as most studies that try to elucidate the underlying transcriptional changes either do not separate different mammary gland cell types or focus exclusively on epithelial cells [20,31,32], the contribution of adipocytes and other stromal cells has been largely overlooked so far. Here we perform genome-wide gene expression analysis of epithelial and stromal mammary cells isolated from wildtype FVB/N mice, either during puberty or at the different stages of the adult estrous cycle. We observe complex but robust patterns of *Wnt* gene expression in basal, luminal, fat and stromal cells. In addition, we detect a cell proliferation signature in basal epithelial cells. The most prominent gene expression changes during the estrous cycle occur in non-epithelial cells, however. These data and findings complement existing studies.

## Results

### Isolation of pubertal and adult estrous-staged mammary glands

To investigate gene expression in specific cell populations of pubertal and adult estrous-staged mammary glands, we wanted to isolate mammary gland tissue at defined time points. The stage of the estrous cycle can be determined by collecting vaginal swabs and scoring the relative proportion of nucleated epithelial cells, cornified epithelial cells and leukocytes in the sample (**Fig. 1a-b**) [14,33–35]. In total, we tracked the estrous cycle of 32 adult (14-16 weeks old) female FVB/N mice every day for at least 10 days (i.e. more than one cycle) to ensure that the mice selected for analysis were showing regular progression through the estrous cycle (examples in **Fig. 1c**). When a smear did not allow us to assign a clear estrous cycle stage, it was scored as a transition smear (e.g. proestrus to estrus), also considering the smear of the preceding day for that particular mouse. To confirm the interpretation of the vaginal smears and increase confidence of our estrous cycle scoring, we also evaluated H&E stained histological sections of vaginal tissue isolated at the time of sacrifice (**Fig. 1d**), since the vaginal epithelium also shows different characteristics at each of the four stages [36,37]. Our analysis of the estrous cycle stage by assessing the tissue sections corresponded with our staging based on vaginal smears (see **Supplementary Fig. 2** for a comprehensive overview of the matching cytology and histology images for every mouse that showed a regular estrous cycle). Of the 32 tracked mice, 7 showed an irregular cycle and were therefore excluded from further analysis (examples in **Fig. 1e**).

**Fig. 1.**
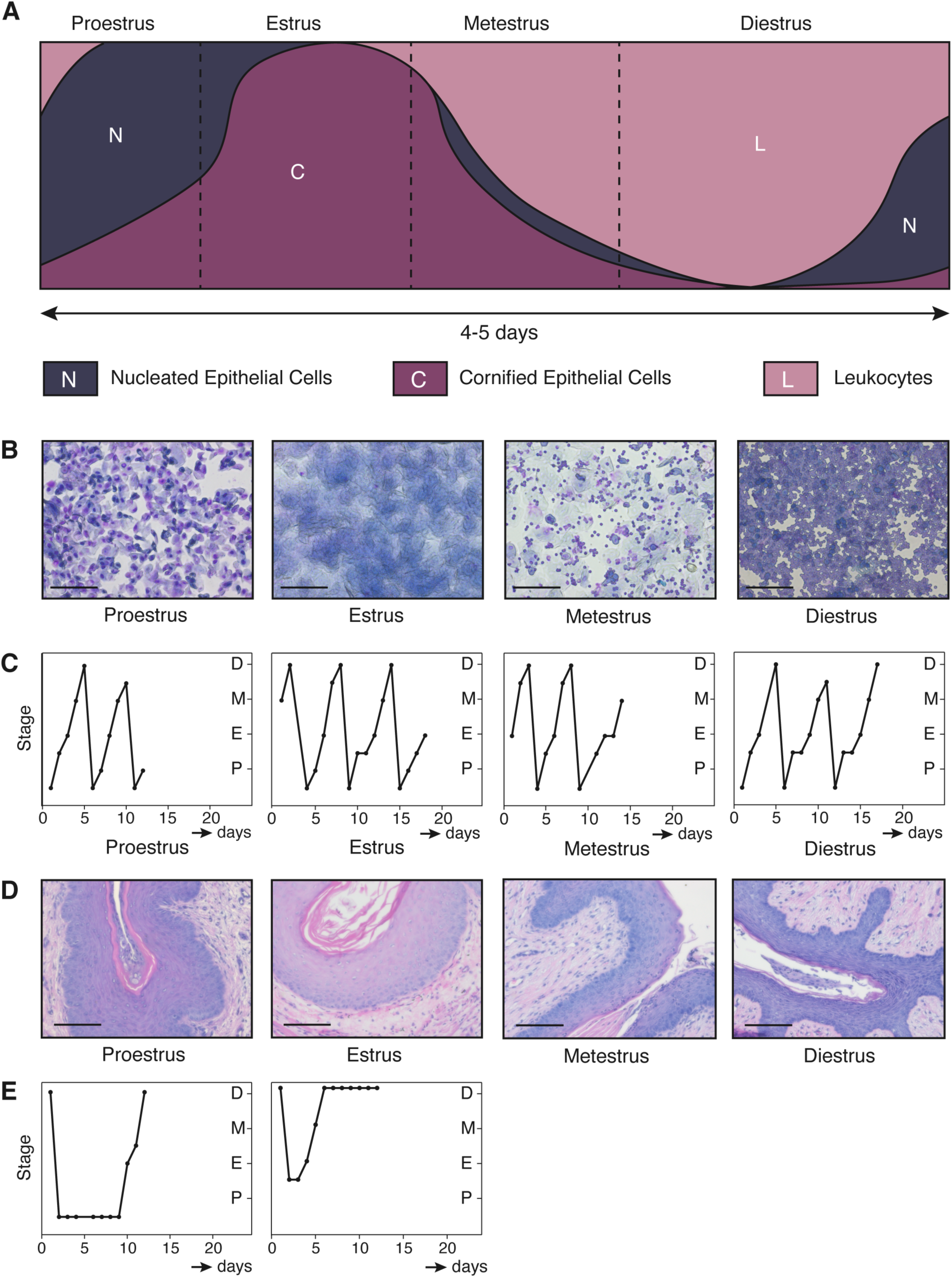
Determining the mouse estrous cycle stage by vaginal cytology. A) Cartoon depicting the relative abundance of cell types present, based on descriptions from the literature [33,34]. Horizontal axis: estimate of the duration of each stage, with the total estrous cycle taking 4-5 days. Dashed lines represent transition into the next stage. B) Brightfield microscopy images of vaginal swabs after staining with Giemsa. Images are from the day of mammary gland isolation for the mice depicted in C). Scale bar = 100 µm. In proestrus, mostly nucleated epithelial cells are present, together with cornified epithelial cells. A small number of leukocytes may be detected in early proestrus stage. When the female mouse is in estrus, mostly cornified epithelial cells are present, together with a small number of nucleated cells (in early estrus) or some leukocytes (in late estrus). Metestrus presents as a mixture of cornified epithelial cells, nucleated epithelial cells and leukocytes. Swabs containing mostly leukocytes indicate that the mouse is in diestrus. C) Graphs depicting monitoring of the estrous cycle. One example is depicted for each endpoint stage, corresponding to the mice in B). D) Brightfield microscopy images of H&E stained formalin fixed paraffin-embedded (FFPE) tissue sections of the vaginal epithelium at different stages of the estrous cycle. One example is depicted for each endpoint stage, corresponding to the mice in B-C). Proestrus can be recognized by a relatively thick epithelial layer and an outer layer consisting of nucleated cells. In estrus the epithelial layer is at its thickest and the outer later is now comprised of cornified cells. Metestrus is characterized by a thinner epithelium, the loss of the cornified cell layers and the transepithelial migration of leukocytes. Diestrus samples show the thinnest layer of epithelium, solely containing non-cornified epithelial cells. E) Two examples of mice with irregular estrous cycles. These mice were excluded from further analysis.

For the adult stages, we ultimately isolated mammary glands from 5 adult female mice in proestrus, 5 in estrus, 4 in metestrus and 4 in diestrus. For each animal, the 3^rd^ and 4^th^ mammary glands were isolated, pooled and processed. For the puberty stage, we selected female FVB/N mice of 34-36 days old (P34-P36, where P stands for postnatal day), since the mammary epithelium is actively undergoing branching morphogenesis at this time point [38]. For each day, the 3^rd^ and 4^th^ glands of 4 individual mice from 3 different litters were isolated, pooled per mouse and processed.

### Number and proportion of cell types remains similar throughout the estrous cycle

To profile gene expression in different cell types, we processed all samples for further analysis, taking care to separate luminal, basal, fat and other stromal mammary cells. Following tissue digestion, we first isolated the adipose fraction of the fat pad (hereafter referred to as adipocytes) by centrifugation. The remainder of the sample was processed to single cells. Using Fluorescence Activated Cell Sorting (FACS) we isolated luminal (lin^-^/EpCAM^high^/CD49f^med^), basal (lin^-^ /EpCAM^med^/CD49f^high^) and stromal (lin-/EpCAM^-^/CD49f^-^) cells (**Fig. 2a**)[39]. **Supplementary Fig. 3** shows the complete gating strategy, as well as a direct comparison of our EpCAM/CD49f staining with the often-used CD24/CD29 staining. We did not measure consistent differences in epithelial cell numbers between different estrous cycle stages (**Fig. 2b**). Furthermore, the percentages of basal, luminal and stromal cells were comparable across different stages (**Fig. 2c**). To confirm this observation, we visualized the morphology of epithelial ducts in mammary glands of stably cycling mice by carmine staining. As we had used all the available tissue for digestion and FACS sorting, we monitored the estrous cycle of a second cohort of FVB/N mice using vaginal cytology (**Supplementary Fig. 4**). A total of 6 mice were used for this analysis, of which 3 were sacrificed in estrus and 3 in diestrus. No clear difference was visible in epithelial morphology between the two stages (**Supplementary Fig. 5**). From these same mice, we used another gland for RNA isolation and qRT-PCR. Expression levels of the progesterone receptor (*Pgr*) were increased in diestrus compared to estrus (**Fig. 3**). This fits with earlier studies [10,14,40] and confirms that the mice were cycling. In contrast, *Wnt4* expression levels between estrus and diestrus were similar, indicating that any changes in the proposed progesterone-mediated induction of *Wnt4* expression are subtle at best [10,20,21,41] and below the detection limit of our qRT-PCR analysis when using whole mammary gland RNA as input. Summarizing, expansion of the epithelial network is not detected at the tissue-wide level (**Fig. 2** and **Fig. 3**) during every estrous cycle in every mouse, in line with a previous report [12].

**Fig. 2.**
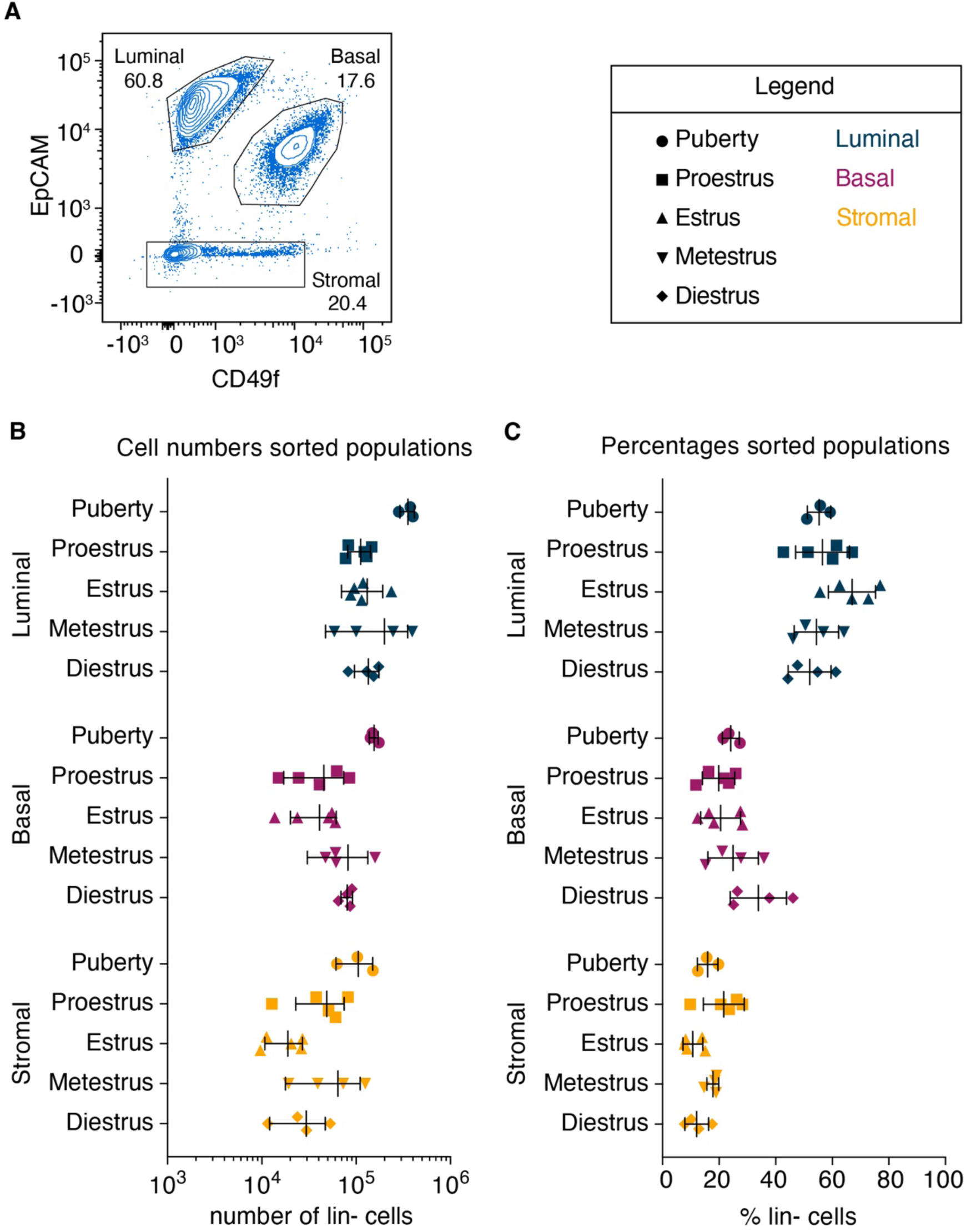
Number and proportion of cell types remains similar throughout the estrous cycle. A) FACS plot depicting the gating strategy for luminal, basal and stromal cells using EpCAM-PE/CD49f-FITC staining. This example is from an adult virgin in proestrus. B) Absolute cell numbers of the sorted populations. Note that the numbers for pubertal and adult mice cannot be compared directly, as for every pubertal replicate sample 4 mice were pooled, whereas all adult mammary glands were sorted individually. Error bars represent mean with SD. C) Same samples as B), but here the relative proportions of the different cell types are plotted. Lin-cells = lineage negative cells.

**Fig. 3.**
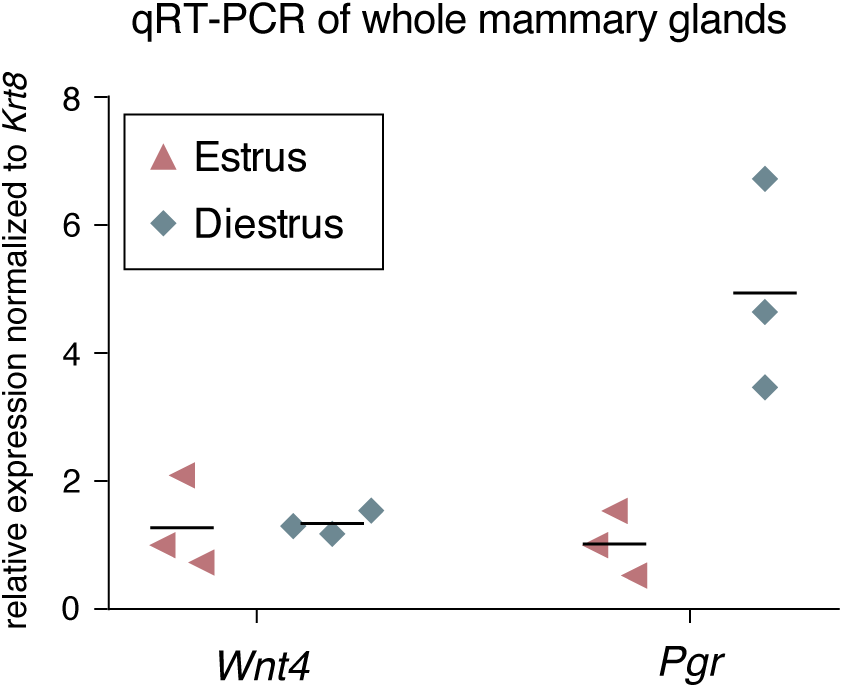
Whole mammary glands in estrus and diestrus show similar *Wnt4* expression. Graph showing *Wnt4* and *Pgr* mRNA expression levels, measured by qRT-PCR on whole mammary gland samples (n=3 mice in estrus and n=3 mice in diestrus). Experimental samples (#4 mammary glands) are from the same mice as those depicted in A. Ct values of *Wnt4* and *Pgr* were normalized to *Krt8* expression, which is a marker for luminal cells. Horizontal lines represent the mean.

### Gene expression changes during the estrous cycle mainly occur in non-epithelial cells

For an unbiased overview of changes in gene expression across different developmental time points and cell types, we performed genome wide expression analyses. Samples from multiple pubertal or multiple adult mice were pooled to obtain sufficient RNA for bulk RNA-seq analysis (see methods for details on how samples were pooled). Two independent RNA sequencing runs were performed, one for the puberty and one for the adult samples. To measure the level of similarity between samples we conducted a multidimensional scaling (MDS) analysis (**Fig. 4a**). As expected, the different samples clustered by cell type and epithelial and non-epithelial cells are separated in the first dimension. With the exception of pubertal basal cells, the two replicates from corresponding developmental stages (i.e. puberty or adult samples) did not consistently cluster together within each cell type. Thus, MDS analysis reveals major differences in gene expression between mammary gland cell types, but not between the selected developmental or estrous cycle stages. Expression analysis of individual cell-type specific markers (luminal: *Krt18* [42]), basal: *Krt14* [42], adipocyte: *Adipoq* [43], stromal: *Col1a1* [44]) (**Fig. 4b**) and hierarchical clustering (**Supplementary Fig. 6**) confirms the correct isolation of defined cell populations.

**Fig. 4.**
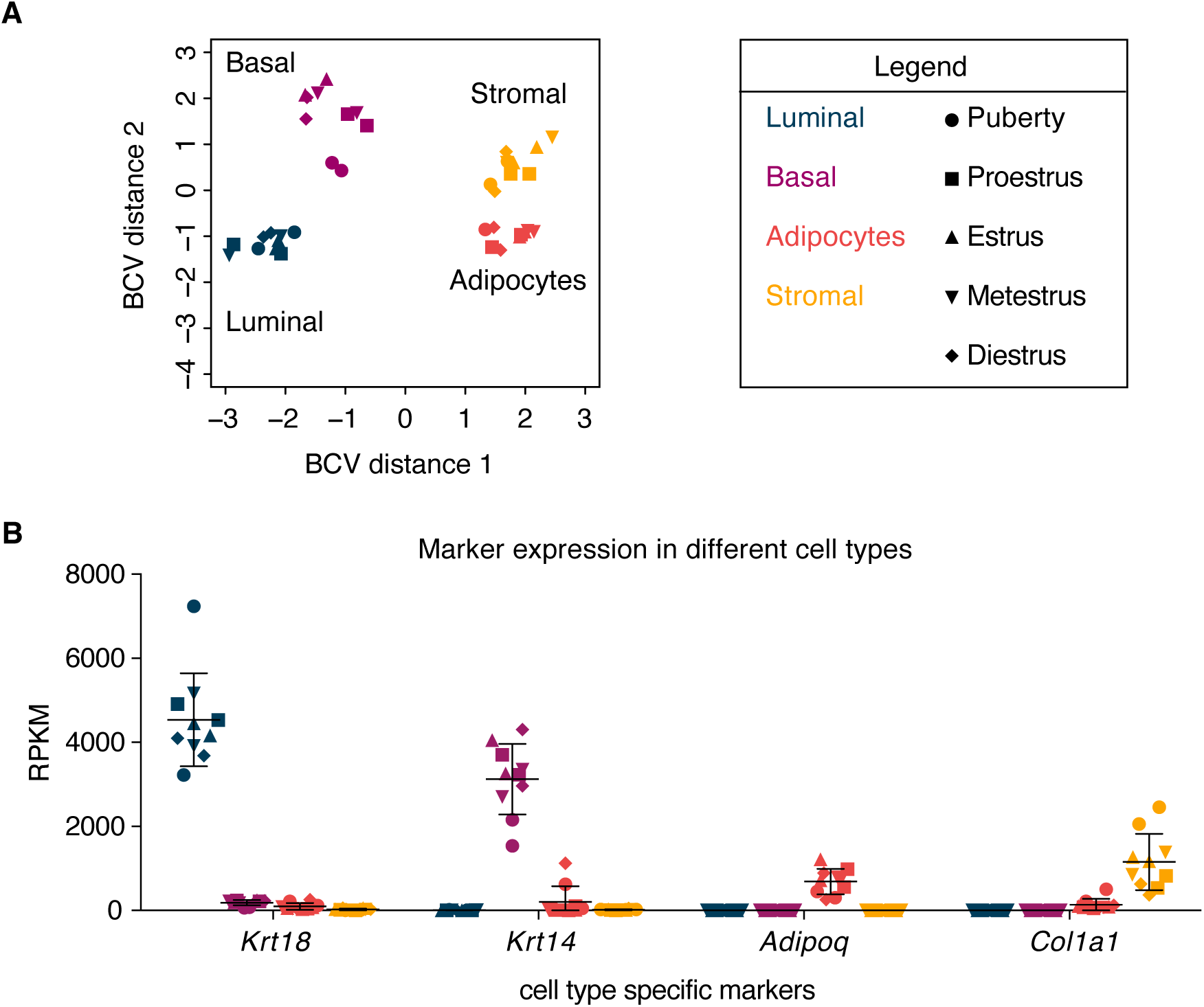
Clustering of different mammary gland cell populations. A) MDS plot of all puberty and adult RNA-seq samples. Distances on the plot correspond to the biological coefficient of variation (BCV) between samples. B) Plot depicting expression of cell type specific marker genes in every sample as Reads Per Kilobase per Million mapped reads (RPKM). Error bars represent mean with SD. In both A) and B) colors represent different cell types and shapes represent different developmental stages. Source data for B) available via https://osf.io/xv83g/ (rpkm_values_allsamples.xlsx).

To compare the different stages of the estrous cycle, we performed differential gene expression analysis (**Fig. 5a**). The luminal cell population, which contains the estrogen- and progesterone-receptor positive cells [45], showed the smallest number of differentially expressed genes in pairwise comparisons of two different estrous cycle stages (21 unique genes in total, FDR<0.05). A total of 255 unique genes were differentially expressed in basal cells, mainly in pairwise comparisons involving the estrus stage (**Fig. 5a**, FDR<0.05). Statistically significant gene expression changes in adipocytes and stromal cells were more prominent. In adipocytes, all pairwise comparisons involving diestrus showed >1000 upregulated genes (**Fig. 5a**, FDR<0.05). In stromal cells, the estrus stage stood out the most, with most differentially expressed genes being downregulated. These results suggest that the non-epithelial cell populations are also hormone responsive, either directly or indirectly, and that reciprocal signaling likely occurs between the different cellular compartments.

**Fig. 5.**
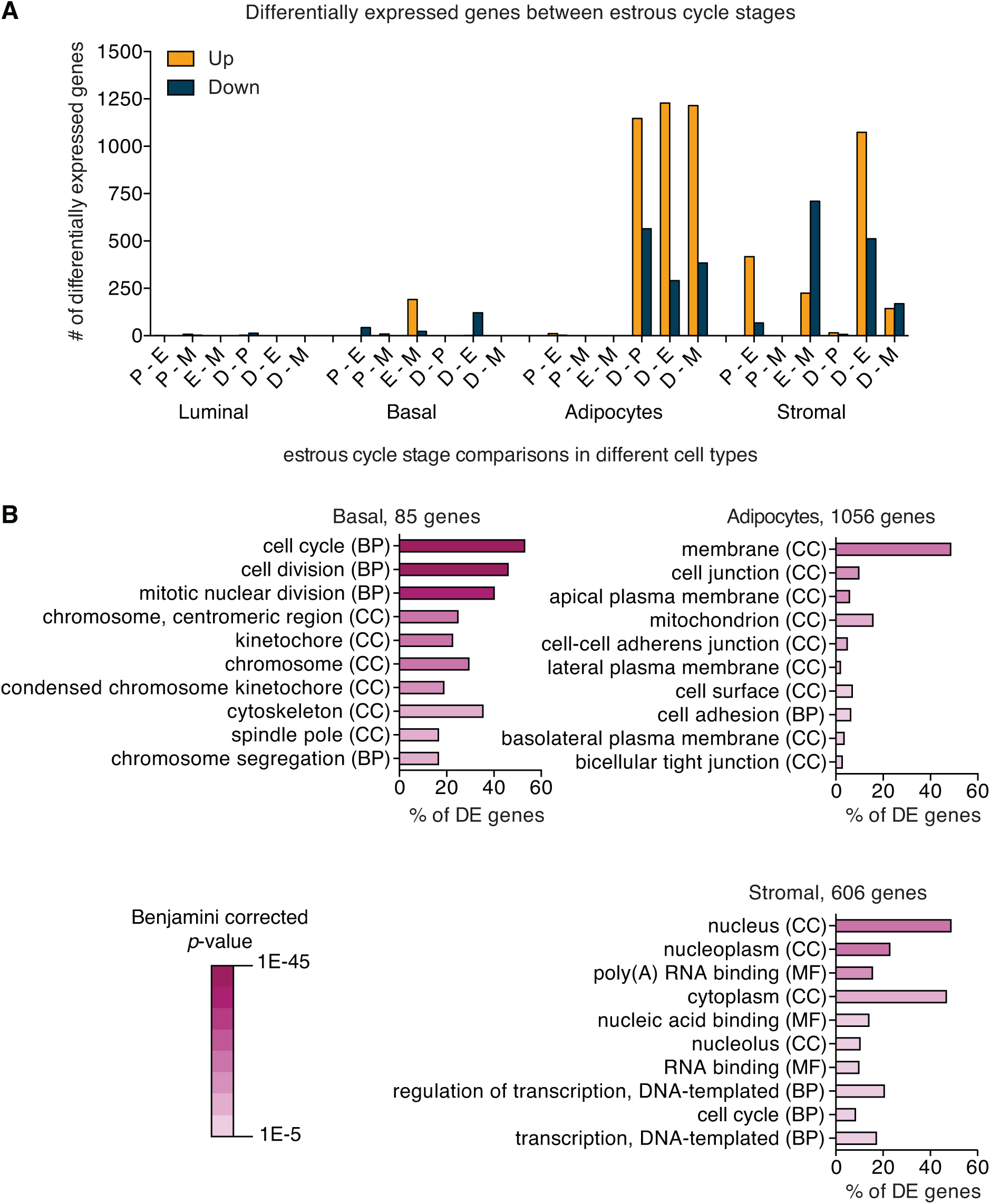
Estrous-cycle dependent changes in gene expression are most prominent in adipocytes and stromal cells. A) Graph depicting the number of differentially expressed genes for each pairwise comparison between different stages of the estrous cycle (FDR <0.05). Yellow bars: upregulated genes. Blue bars: downregulated genes. On the X-axis are different pairwise comparisons of different estrous cycle stages. P = proestrus, E = estrus, M = metestrus, D = diestrus. B) Graphs depicting functional enrichment analysis of all differentially expressed genes (DE genes) in basal cells, adipocytes and stromal cells (FDR <0.01) in DAVID (version 6.8)[48,49]. Per cell type the top 10 most enriched GO terms are shown. Color intensity indicates Benjamini corrected p-value. CC = cellular component, BP = biological process, MF = molecular function. Source data available via https://osf.io/xv83g/ (Fig5A_sourcedata_numbers.xlsx, Fig5A_sourcedata_FDR005 and Fig5B_sourcedata_FDR001).

Functional gene ontology (GO) classification of differentially expressed genes (cut off FDR<0.01, **Fig. 5b**) [46,47] revealed that differentially expressed genes in adipocytes are mostly linked to *cell membrane* and *cell adhesion*, while the differentially expressed gene list in stromal cells is enriched for genes associated with *nucleus* and *regulation of transcription*. The most prominent enrichment for specific GO terms was observed in basal cells and is related to *cell cycle*/*cell division* (see **Supplementary Table 1** for a selection of differentially expressed genes with the highest fold change (logFC of >|5|) in basal cells and a logCPM of >1). A cell cycle gene signature containing 65 out of the total 85 genes that are differentially expressed in basal cells (FDR<0.01) showed elevated expression in basal and luminal cells, but not in non-epithelial cells in estrus compared to other estrous cycle stages (**Supplementary Fig. 7**). However, this does not translate to clear changes in the number of cells after sorting specific cell types during at diestrus (**Fig. 2**), nor in readily apparent side-branching (**Supplementary Fig. 5**).

### A complex *Wnt* gene expression landscape in the mammary gland

Because the expression of *Wnt4* and other *Wnt* genes has been linked to steroid hormones in different tissues [10,20,21,50–53], we zoomed in on the expression of individual *Wnt* genes. Neither basal nor luminal epithelial cell types showed differential expression of any *Wnt* gene across the estrous cycle. We did, however, observe differential expression of 6 *Wnt* genes in non-epithelial cells (**Supplementary Table 2**, cut off: FDR <0.05 in pairwise stage comparisons). In adipocytes, *Wnt4, Wnt5b, Wnt7b* and *Wnt10a* are upregulated in diestrus. In stromal cells, *Wnt6* and *Wnt5a* are downregulated in diestrus compared to estrus and metestrus, respectively. We do point out that the absolute expression levels of these *Wnts* in stromal cells and adipocytes are often close to background levels (**Fig. 6**). Note that *Wnt* expression itself is not inherently low: Multiple *Wnt* genes are expressed at decent levels in luminal and basal cells – and although *Wnt4* does not meet our cut off for statistical significance, we do detect an ∼1.7 fold difference in luminal *Wnt4* expression between proestrus (mean RPKM:17.2) and diestrus (mean RPKM: 10.1, **Fig. 6**).

**Fig. 6.**
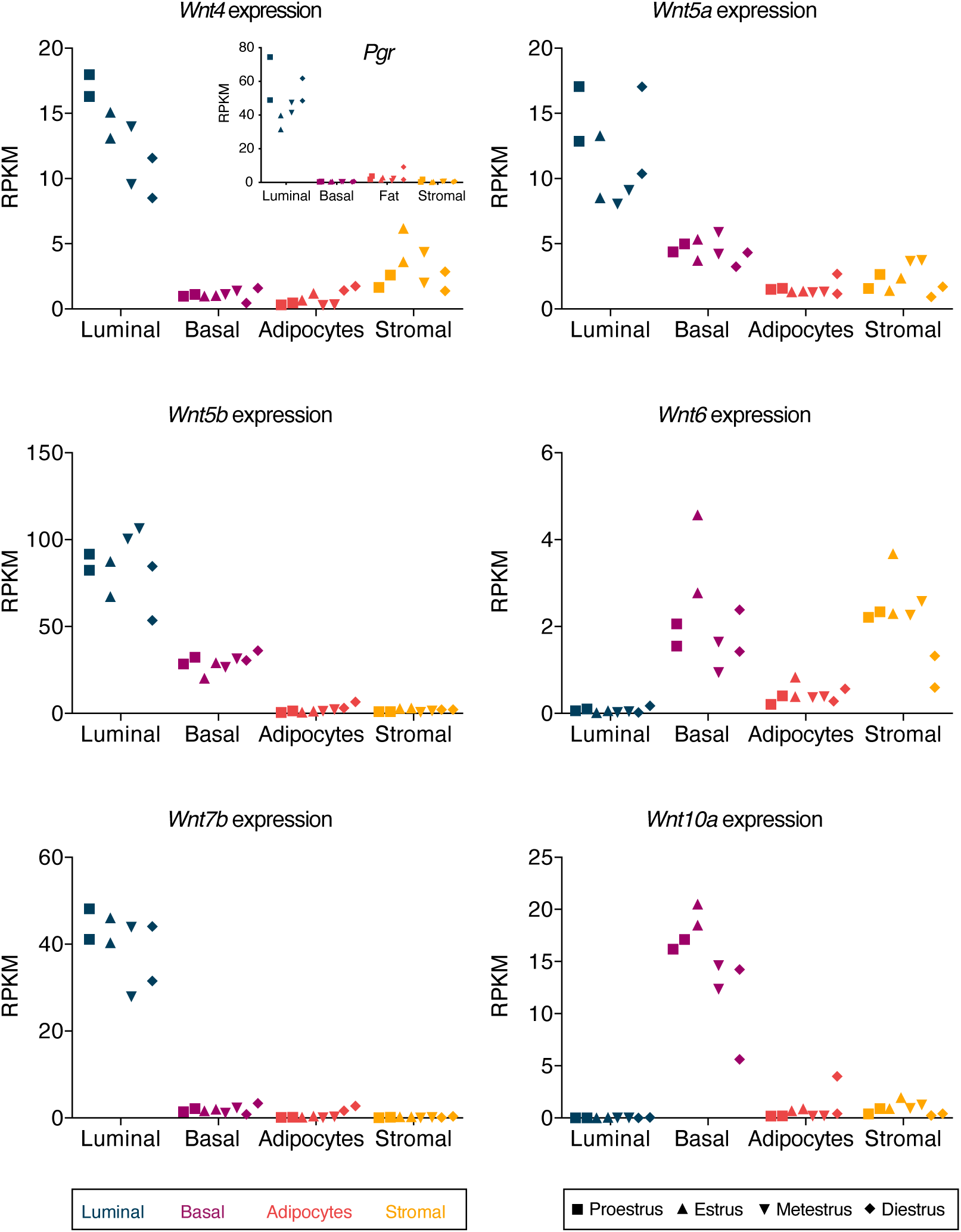
Expression levels of differentially expressed *Wnt* genes in different mammary gland cell types during the estrous cycle. RNA-sequencing results from different mammary gland cell types and different stages of the estrous cycle. Dot plots depict the expression of 6 individual *Wnt* genes that were found to be differentially expressed (FDR <0.05) in pairwise comparisons of different estrous cycle stages (Table 3). RPKM = Reads Per Kilobase per Million mapped reads. Inset in the top left panel shows *Pgr* expression, which rises and peaks prior to *Wnt4* expression. Colors represent different cell types and symbols represent different stages of the estrous cycle. Source data available via https://osf.io/xv83g/ (rpkm_values_allsamples.xlsx).

Finally, we performed unsupervised hierarchical clustering of our pubertal and estrous cycle staged samples (**Fig. 7**). This revealed two interesting features. First, epithelial and non-epithelial mammary cell populations cluster according to cell type based solely on the expression of different *Wnt* genes. Second, the expression patterns of individual *Wnt* genes vary between cell type rather than developmental time point. In puberty, *Wnt4* and *Wnt7b* expression is restricted to luminal cells, while *Wnt6* and *Wnt10a* expression is restricted to basal cells. *Wnt5a* and *Wnt5b* are expressed in both luminal and basal epithelial cells. This is in line with earlier work that showed expression of these genes in the pubertal mammary epithelium using microarray gene expression analysis [54]. *Wnt4*, *Wnt5a*, *Wnt5b* and *Wnt7b* remain expressed in the adult luminal cell population. This fits with published microarray and transcriptomic data [54–56], and with studies using RNA *in situ* hybridization or qRT-PCR analysis [20,21,26,57,58]. In the epithelium, expression of *Wnt6* and *Wnt10a* remains restricted to the adult basal layer, also corresponding to previous studies [55,56]. *Wnt5a*, *Wnt5b*, *Wnt9a*, *Wnt10a* and *Wnt11* are also expressed in basal cells, in line with published scRNA data [56]. Although expression of *Wnt2* and *Wnt6* in stromal cells has been reported [58,59], most genome wide expression analyses have so far focused on mammary epithelial cells. We find that *Wnt2* is indeed mainly expressed in stromal cells, in agreement with RNA *in situ* hybridization experiments [60]. In adipocytes, *Wnt* gene expression was relatively low, although these cells showed some expression of *Wnt2*, *Wnt10b*, and *Wnt16*. Stromal cells express multiple *Wnt* genes in addition to *Wnt2*, including some *Wnt4* and *Wnt6* (**Fig. 6**), with only *Wnt9b* and to a lesser extent *Wnt2b* expression being exclusive for the stromal cell population. The precise regulation and function of these *Wnt* gene expression patterns warrant further research.

**Fig. 7.**
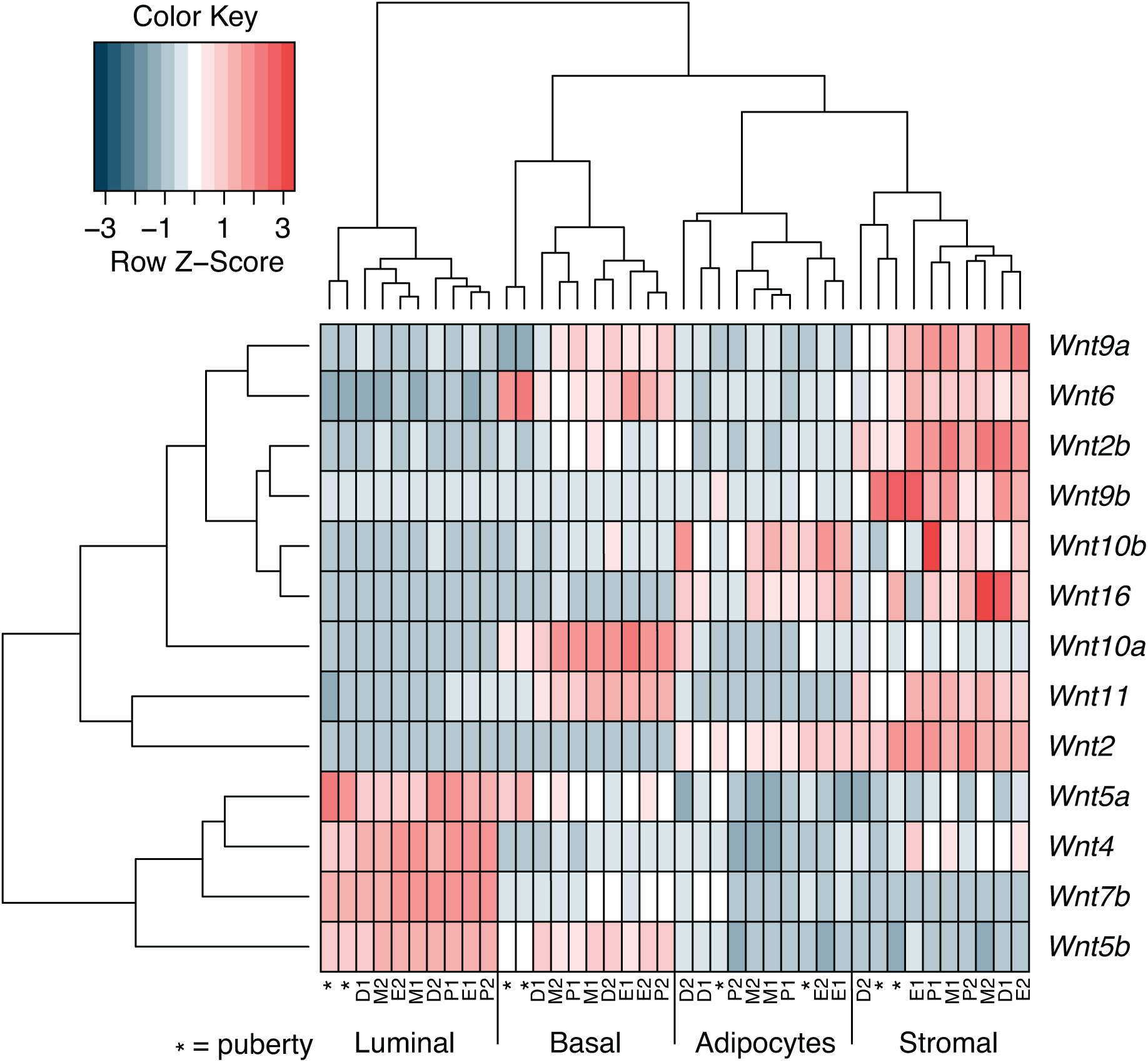
RNA-sequencing reveals complex spatiotemporal expression patterns of individual Wnt genes. Unsupervised hierarchical clustering of *Wnt* gene expression in different mammary gland cell types at different developmental time points. Labels for different samples: * = puberty samples, P = proestrus, E = estrus, M = metestrus, D = diestrus. Numbers “1” and “2” indicate replicate samples (see methods for details). RPKM = Reads Per Kilobase per Million mapped reads. Values were normalized for each gene across all samples using Z-scores. Source data available via https://osf.io/xv83g/ (rpkm_values_allsamples.xlsx).

## Discussion

Few studies to date have sampled the basal, luminal, adipocyte and stromal cell populations in pubertal and adult mice. The specific aim of this study was to better understand the transcriptional response of the epithelial and non-epithelial mammary cell populations to physiologically changing hormone levels, with a specific focus on estrous-cycle staging. Given the role of WNT signaling in mammary gland development and tissue homeostasis, including a prior link to mammary stem cell maintenance and steroid hormone responsiveness, we paid specific attention to *Wnt* gene expression patterns and changes therein.

### Determining estrous cycle stage by vaginal cytology

Different methods exist for assessing the stage of the estrous cycle in adult female mice. While it can be done by assessing the overall appearance of the vaginal opening [61], metestrus and diestrus are difficult to distinguish using only visual inspection. It is therefore recommended to assess the ratio of different cell types using vaginal smears [34]. For this study, we confirmed our stage determination by vaginal cytology with histological analysis of the vaginal epithelium (**Supplementary Fig. 2**). While sampling our cohort, we found that 7/32 mice were not cycling reliably, despite being housed in the vicinity of male mice in the same open cages as their cycling littermates (**Fig. 1e**). This highlights the importance of monitoring the cycle for multiple days in a row: only collecting swabs on the day of tissue isolation or only collecting samples once or twice a week is insufficient to determine the precise stage and progression of the estrous cycle [32]. As shown previously, proper estrous cycle stages requires samples to be collected every day, at the same time, for at least one week [10,11,13,62]. Indeed, daily monitoring for at least 8 days allowed us to identify animals with irregular cycles and exclude these mice from further analysis.

### Epithelial outgrowth does not occur in every mouse in every cycle

Studies investigating morphological changes of the mammary epithelium during the estrous cycle have reported conflicting results [7,8,10–13,16]. Although findings concerning the timing of apoptosis are consistent (diestrus [7,10,12,13,16]), it remains unclear whether side branching occurs in proestrus/estrus [7,12,16] or diestrus [10,11,13]. Assuming that outgrowth of the epithelium requires cell division, one would expect a burst in cell proliferation to precede the appearance of tertiary branches. Our RNAseq data show a cell division/mitosis signature in basal cells at estrous (**Fig. 5**, **Supplementary Table 1** and **Supplementary Fig. 7**), suggesting cell proliferation or a mitotic arrest, but we did not observe accompanying or subsequent changes in cell numbers (**Fig. 2**) or morphology (**Supplementary Fig. 5**) across the stages of the estrous cycle. Tissue level analyses at single cell resolution, including intravital imaging approaches [63], will be required to understand how, when and where fluctuations in systemic hormone levels are translated into a locally controlled cell division, differentiation and migration response. Altogether, we and others [12] find that proliferation of epithelial cells during the estrous cycle is not as black and white as reported in some studies. Rather, proliferative heterogeneity exists. From our set up and analysis we cannot exclude confounding factors such as environmental conditions (e.g. housing and diet) or effects arising from sampling different anatomical locations (e.g. #3 versus #4 mammary gland). It is likely, however, that true differences exist between strains, between individual animals, and possibly between different regions of the tissue. Our findings suggest that cell proliferation is a local, rather than a global event, in the same way that some terminal end buds (TEBs) during puberty are more proliferative than others [64].

### Gene expression changes between different estrous cycle stages

We did not detect robust estrous-cycle dependent changes in gene expression, but perhaps we cannot expect major fluctuations in gene expression when we do not clearly observe the expected corresponding morphological changes (**Fig. 2**, **Supplementary Fig. 5**). We also may have missed differences due to our experimental setup. First, we isolated the bulk luminal cell population. As such, the effect of fluctuating steroid hormone levels might be masked, since the estrogen and progesterone receptor (ER and PR) are only expressed in a subset of luminal cells in adult mice [45]. Either enriching for ER+ and PR+ mature luminal cells via FACS sorting [65–67], or single-cell rather than bulk RNAseq [56], would facilitate gene expression analysis of hormone receptor expressing cells specifically. Second, while we harvested glands from 4-5 mice for each stage, we had to pool isolates to obtain sufficient RNA. This unfortunately reduced the number of biological replicates in our analysis. Third, it is also possible that rapid changes in morphology as previously observed, are not caused by changes in gene expression. In fact, other mechanisms may enable faster and/or more dynamic changes in cell behavior. This includes alternative splicing, which several studies have shown to be important in vertebrate tissue development and homeostasis [68,69]. Changes to the poly(A) tail of mRNA can also play a regulatory role by changing translation efficiency, mRNA stability and degradation [70]. Post translational modifications such as phosphorylation, acetylation or ubiquitination can change protein functionality and add another level of regulation [71].

### Non-epithelial cells should be included in studies of mammary gland biology

Pairwise comparisons of the different estrous cycle stages revealed that most differentially expressed genes were found in non-epithelial rather than in epithelial cell populations (**Fig. 5a**). While hormone receptor positive cells are indeed also found outside of the mammary epithelium [31,72], the difference in absolute numbers of differentially expressed genes between the epithelial and non-epithelial mammary cell types, is remarkable. This adds to the evidence that molecular cues that control epithelial cell behavior during the estrous cycle are not only coming from the epithelial cells itself, but also from adipocytes and stromal cells [73]-either in direct response to hormones or in response to hormone-induced paracrine signals.

Some caution is warranted when interpreting these results. First, we isolated adipocytes via differential centrifugation after the initial enzymatic digestion step, possibly resulting in less pure samples than for the other cell types, which were isolated by FACS sorting. While our quality controls do not show any evidence of impurity (**Fig. 4** and **Supplementary Fig. 5**), we cannot exclude some contamination. Second, the FACS sorted stromal population is not well defined. While we made sure to exclude doublets, dead cells, hematopoietic and endothelial cells (**Supplementary Fig. 3**), our sorting of mammary stromal cells was not based on the presence of specific stromal markers, but rather on the absence of epithelial markers (**Fig. 2a**). Because fibroblasts are the main component of the mammary stroma and we did not actively exclude them in any step prior to sorting, we expect these cells to be the main component of our sorted stromal cell population. This is supported by the specific expression of *Col1a1* in this population (**Fig. 4b**).

### *Wnt* genes show defined spatial gene expression patterns in the mammary gland

We did not detect statistically significant changes in *Wnt4* expression in luminal cells (**Fig. 6**), where it has been reported to act downstream of progesterone [10,20,21,41]. In some of these prior studies, *Wnt4* expression was measured after treating ovariectomized mice with 17ý-estradiol and progesterone. The fact that we measure gene expression under physiological conditions might be one explanation for the discrepancy, since endogenous hormone fluctuations, and their downstream effects, are likely to be much smaller. This is supported by the fact that we did not detect a correlation between *Pgr* and *Wnt4* gene expression in whole mammary gland mRNA (**Fig. 3**), and by the fact that we were able to detect subtle, non-statistically significant changes in *Pgr* (∼1.5 fold difference between diestrus/proestrus versus estrus/metestrus, mean RPKM of 58.4 and 40.1, respectively) that preceded equally subtle changes *Wnt4* expression (∼1.7 fold difference between proestrus and diestrus, mean RPKM of 17.2 and 10.1, respectively) in our FACS sorted, and thus partially enriched, samples (**Fig. 6**).

As far as we know, spatiotemporal *Wnt* gene expression differences across luminal, basal, stromal and adipocyte populations during puberty and across the four stages of the estrous cycle has not been examined before. Previous transcriptomic analyses that distinguished luminal cells based on their hormone receptor expression status [55,56] showed that *Wnt4*, *Wnt5a* and *Wnt7b* are mainly (but not exclusively) expressed in hormone receptor positive luminal cells. We can conclude that multiple *Wnt* genes show well defined spatial gene expression patterns, resulting in a complex *Wnt* gene expression landscape in basal, luminal, adipocyte and stromal cell populations in the postnatal mammary gland.

### Conclusion and outlook

Although a link between the reproductive cycle and breast cancer development is known to exist, the molecular basis of this interaction remains incompletely understood. This work contributes to understanding the effects of physiological hormone fluctuations on gene expression and the molecular mechanisms underlying dynamic morphological changes in the mammary gland during the reproductive cycle. It shows that endogenous gene expression changes in response to physiological changes in steroid hormone levels are subtle. Our dataset should be a useful resource for future studies investigating gene expression patterns in different cell types of the mammary gland across different developmental timepoints. It also informs future studies, which should ideally include a higher number of replicates to capture sample-to-sample heterogeneity and should continue to consider early and late timepoints of each estrous cycle stage, while at the same time isolating hormone-receptor positive and negative cells for both epithelial and non-epithelial cell populations to reduce intra-sample heterogeneity. Ideally, these experiments would be performed at single-cell resolution. Ultimately, a better comprehension of the molecular mechanisms underlying the dynamic and heterogenous changes in mammary cell behavior in response to systemic hormones and paracrine growth factors will increase our understanding of reproductive cycle-related breast cancer risks. In the long run, these insights have the potential to contribute to the prevention and personalized treatment of breast cancer in premenopausal women [74].

## Materials and methods

### Mice

Wildtype, inbred FVB/NHan®Hsd mice (referred to as FVB/N in the main text) were purchased from Envigo. Breeding and estrous cycle monitoring of the experimental cohort was performed in-house. All mice were maintained under standard housing conditions in open cages, with 12 hour light/dark cycle and *ad libitum* access to food and water. Experiments were performed according to institutional and national guidelines and regulations (see study-specific approval). Pubertal mice were 34-36 days old. N=4 mice from n=3 different litters were pooled to obtain sufficient cells for RNA-sequencing according to **Table 1**. Adult mice were 14-16 weeks old and were sacrificed at the appropriate stages of the estrous cycle according to **Supplementary Fig. 2**. Samples from n=2 or n=3 different mice were pooled according to **Table 1** to obtain sufficient material for RNA-sequencing.

**Table 1.** Mammary glands of different mice were pooled per stage.

| Timepoint | Puberty |  |  | Adult |  |  |  |  |  |  |  |
| --- | --- | --- | --- | --- | --- | --- | --- | --- | --- | --- | --- |
|  | P34 | P35 | P36 | P1 | P2 | E1 | E2 | M1 | M2 | D1 | D2 |
| # of mice in pooled sample | 4 | 4 | 4 | 2 | 3 | 2 | 3 | 2 | 2 | 2 | 2 |

For the puberty dataset, samples from n=4 mice were pooled prior to FACS sorting. P34, P35 and P36 represent three consecutive days of mammary gland isolation. P = proestrus, E = estrous, M = metestrus, D = diestrus. For adult timepoints, samples from individual mice were FACS sorted separately and pooled at the time of RNA isolation. For each estrous cycle stage, samples from n=2 or n=3 mice were pooled during RNA isolation, based on our combined interpretation of vaginal cytology and histology (Supplementary Fig. 2). If no clear difference between early (“1”) or late (“2) stages was observed, we used cell number as a second criterium to achieve equal RNA input for next generation sequencing. For the puberty samples, we selected 2 replicates based on RNA quality. For each estrous cycle stage, we ultimately also ended up with 2 replicates, each containing RNA from 2 or 3 mice.

### Vaginal cytology

Vaginal swabs were collected using plastic Pasteur pipettes and PBS. From the tip of the vaginal opening, the vagina was flushed 2-3 times with a few drops of PBS. The sample was transferred to a glass slide, air-dried at 37°C, stained with Giemsa (Sigma-Aldrich cat. #48900) for 30 seconds and rinsed with PBS. Animals of the right stage were sacrificed as soon as possible after the samples had been stained and assessed under the microscope. Brightfield images of the stained cytology samples were taken at 20x magnification using a Zeiss Axio Vert.A1 microscope.

### Histology

Vaginal tissue samples were fixed in 4% PFA for 24 hours, dehydrated through ascending grades of ethanol, cleared in Histo-Clear II (National Diagnostics cat. #HS-200) and embedded in paraffin. Five µm thick tissue sections were cut and mounted on glass slides. Slides were baked at 60°C for 45 minutes, deparaffinized in Histo-Clear II, rehydrated through descending grades of ethanol, stained with 50% Mayer’s Hematoxylin (Sigma-Aldrich) for 30 seconds, rinsed for 5 minutes in tap water, washed in PBS for 3 minutes and 70% ethanol for 5 minutes, stained with Eosin Y (Sigma-Aldrich, cat. #HT110132) for 2 minutes, dehydrated (through 70% ethanol, 100% ethanol and 100% isopropanol), cleared in Histo-Clear II (National Diagnostics cat. #HS-200) and mounted with a coverslip. Brightfield images of the histology samples were taken at 20x magnification using a Zeiss Axio Vert.A1 microscope.

### Mammary gland digestion

The 3^rd^ and 4^th^ mammary glands of FVB/N mice were isolated, manually minced and enzymatically digested in an orbital shaker for 2 hours at 37°C in digestion mix (9.2 ml DMEM/F12, 5% FCS, 1% Penicillin/Streptomycin, 25 mM HEPES (Gibco, cat. #15630056) and 300 U/ml Collagenase IV (Gibco, cat. #17104019), with 10 ml digestion mix per mouse for 4 glands per mouse). After centrifugation, the fat fraction was obtained by pipetting and transferring the top layer to TRIzol LS (Invitrogen, cat. #10296028). Samples were stored at -80°C. The remainder of the sample was processed for single cell isolation. For this, cell pellets were resuspended in HBSS (Gibco, cat. #11540476 supplemented with 2% FBS (Gibco, cat. #11573397) and ACK solution (Gibco, cat. #A1049201) (1:3) and incubated at room temperature (RT) for 5 minutes to lyse red blood cells. To dilute and inactivate the ACK buffer, 13 ml of HBSS was added and cells were spun down for 5 min, 1000 rpm, 4°C, and brake set to 1. Cell pellets were resuspended in 2 ml pre-warmed 0.05% Trypsin-EDTA (Gibco, cat. #11590626) and incubated for 5 minutes at 37°C, after which 3 ml of pre-warmed serum-free DMEM and 1 µg/ml DNAseI was added to the solution. After mixing, 8 ml of DMEM with 10% FCS was added to stop trypsinization. Cells were filtered through a 40 µm mesh into a fresh tube.

### Antibody staining and FACS

Cells were resuspended in 200 µl HBSS supplemented with 10% FBS and a cocktail of the following antibodies: EpCAM-PE (1:100, eBioscience, 12-5791-82, clone G8.8), CD49f-FITC (1:100, eBioscience, 11-0495-82, clone GoH3), CD45-Bio (1:100, eBioscience, 13-0451-82, clone 30-F11), CD31-Bio (1:100, eBioscience, 13-0311-81, clone 390), Ter119-Bio (1:100, eBioscience, 13-5921-81, clone TER-119). After incubating in the dark on ice for 20 minutes, cells were washed twice with HBSS supplemented with 2% FBS and incubated in 200 µl HBSS supplemented with 10% FBS containing Streptavidin-APC (1:200, eBioscience, 17-4317-82). After antibody staining, cells were stained with DAPI (1:5000 (Invitrogen cat. #D1306) and filtered through a 50 μm mesh. Cells were sorted using a BD FACS Aria III. FITC was excited with a 488 nm laser and emission was filtered using a 530/30 nm bandpass filter. PE was measured using a 561 nm laser and 582/15 nm bandpass filter. DAPI was excited with a 407 nm laser and emission was filtered using a 450/50 nm bandpass filter. APC was measured using a 633 nm laser and 660/20 nm bandpass filter. Cells were sorted with a plate voltage of 2500 V using the 4-Way Purity precision mode. Sorted cells were collected in TRIzol LS (Invitrogen, cat. #10296028). Post-sort purity checks were performed after every sort and were always >90%. Samples were stored in TRIzol LS at -80°C.

### Carmine staining

Freshly isolated 3^rd^ mammary glands of estrous cycle monitored mice were flattened between two glass slides, incubated on a nutator at RT for 4 hours in a 50 mL tube containing 12.5 ml 100% EtOH and 12.5 ml acetic acid. Glands were removed from the glass slides, incubated in 70% EtOH for 1 hour on a nutator at RT and rinsed in demi-water and stained overnight (O/N) in carmine solution (1g carmine (Sigma, cat. #C1022), 2.5 g aluminum potassium sulphate (Merck, cat. #101047), 500 ml water, boiled for 20 minutes and filtered). After staining, glands were washed with 100% EtOH for 4 hours. Glands were cleared and stored in Histo-Clear II at RT. Images were taken on a Leica stereomicroscope at 5x magnification.

### qRT-PCR

Total RNA from the 4^th^ mammary glands was isolated from TRIzol (Invitrogen, cat. #15596018) according to manufacturer’s guidelines. After DNAse treatment, cDNA synthesis was performed from 4 µg RNA using SuperScript IV Reverse Transcriptase (Invitrogen, cat. #18090200) and Random Hexamers (Invitrogen, cat. #N8080127) according to manufacturer’s guidelines. During the reverse transcriptase reaction, RiboLock RNase Inhibitor (Thermo Scientific, cat. #EO0328) was added. After completion of cDNA synthesis, samples were diluted 10x and qRT-PCR reactions were performed using a QuantStudio 3 Real-Time PCR System (Applied Biosystems). For each reaction, 5 µl of diluted cDNA was added to a mix of 4 µl 5X HOT FIREPol EvaGreen qPCR Mix Plus (ROX) (Solis Biodyne, cat. #08-24-00008), 1 µl primers (0.5 µl forward and 0.5 µl reverse, from a 10 µM stock) and 10 µl nuclease-free water. Reactions were performed in triplicate in a 96×0.2 ml plate (BIOplastics, cat. #AB17500). Thermal cycling reactions included the following stages: 2 min. at 50.0°C and 15 min. at 95.0°C, then 40 cycles of 15 sec. at 95.0°C and 1 min. at 60.0°C, followed by the melting curve stage. The following primers were used: *Wnt4* forward: ACTGGACTCCCTCCCTGTCT, *Wnt4* reverse: TGCCCTTGTCACTGCAAA, *Pgr* forward: TGCACCTGATCTAATCCTAAATGA, *Pgr* reverse: GGTAAGGCACAGCGAGTAGAA, *Krt8* forward: AGTTCGCCTCCTTCATTGAC, *Krt8* reverse: GCTGCAACAGGCTCCACT,

### RNA-sequencing

RNA extraction, purification and sequencing, as well as data processing until read count calculation was done at the NKI Genomics Core facility. For the puberty samples (see **Table 1**), the following samples were used based on quality control: Replicate 1: Basal P34, Luminal P34, Stromal P34, Fat P34; Replicate 2: Basal P35, Luminal P35, Stromal P35, Fat P36. Adult samples were pooled according to **Table 1**. RNA was extracted from TRIzol LS and purified using the Qiagen RNeasy column purification kit. RNA quality was checked with a Bioanalyzer (Agilent), after which polyA+ stranded RNA library preparation was performed using the Illumina TruSeq stranded RNA prep kit. RNA-sequencing was performed on a HiSeq 2500 (Illumina) System at the NKI Genomics Core Facility.

### Bioinformatics

Single-end reads (65 bp) were aligned to reference genome GRCm38/mm10 with Tophat version 2.1 and Bowtie version 1.1.0 [75]. Expression values were determined by HTSeq-count [76]. Raw gene-level count tables were processed further using edgeR (version 3.28.0) [77,78] and limma (version 3.42.1) [79]packages with R (version 3.6.2) [80]. A cutoff of CPM > 1 in at least 2 libraries was applied to filter out genes with low counts prior to trimmed mean of M-values (TMM) normalization. To visualize distances between gene expression profiles based on the biological coefficient of variation, the plotMDS function was used with method set to “bcv”. RPKM values were generated with the RPKM function after calculating exonic region lengths from non-overlapping exons per transcript ID (genome version mm10). The edgeR glmQLFit function was used to estimate the dispersion trend before testing for differential expression between different estrous cycle stages. Unless otherwise noted, a false discovery rate (FDR) of 5% was considered as a threshold for significance. The heatmap of *Wnt* gene expression in different mammary gland cell types was created with the heatmap.2 function (gplots) with Z-score scaling along rows and dendrograms computed using default settings. The heatmaps for Supplementary Fig. 5 and 6 were made with the pheatmap package (version 1.0.12) [81].

### Software and online resources

Differential gene expression analysis for Figures 6-9 was performed in R and R Studio as described under Bioinformatics. Gene set enrichment analysis for Figure 7 and Supplementary Figure 7 was performed using DAVID (https://david.ncifcrf.gov/ [82]) and MSigDB (https://www.gsea-msigdb.org/gsea/msigdb [83,84]. Plots and graphs were made in R Studio and Graphpad Prism. Figure 1 was drawn in Adobe Illustrator. All figures were assembled in Adobe Illustrator. Tables were made in Microscoft Word. Spreadsheets were made in Microscoft Excel.

### Data availability

RNA-seq source data for puberty and adult samples have been uploaded to NCBI GEO and can be accessed via GSE213451 as raw counts, with links to the SRA read archive to access the original .fastq files. Source data for the figures and tables are provided via the Open Science Framework and can be accessed through https://osf.io/xv83g/ (identifier: DOI 10.17605/OSF.IO/XV83G). This includes processed RNAseq data (provided as RPKM values in both .txt and .xlsx format), as well as spreadsheets with source data for **Fig. 5**, **Supplementary Fig. 1, 6 and 7** and **Supplementary Table 1**.

### Figures and Figure Captions

For this initial submission all figures and figure captions have been provided in line with the text.

This manuscript has 7 Supplementary Figures, which are appended below the reference section. For all other source data, see ‘Data availability’ in the Materials and Methods section.

## Statements and Declarations

### Study-specific approval

All animal experiments were approved by the Animal Welfare Committee of the University of Amsterdam (2013-2017: local permit) or the Centrale Commissie Dierproeven (2018-2022: AVD1110020173145).

## Acknowledgements

We thank our animal caretakers for taking daily care of the mice and the NKI Genomics Core facility for performing the RNA-sequencing experiment, including the mapping and initial quality control of the data. We acknowledge Amber Zeeman for training and support with the estrous cycle staging. We thank our colleagues from the section of Molecular Cytology at the Swammerdam Institute for Life Sciences for discussions and feedback during the project and Thijs van Boxtel for critical reading of the final manuscript.

## Author contributions

Conceptualization: KEW, NH and RvA conceived the study; Data curation: KEW, NH and RvA were responsible for research data management; Formal analysis: KEW, NH and RvA performed data analysis; Funding acquisition: JJ, KEW, RvA; Investigation: KEW and NH designed and performed all experiments (NH and KEW: monitoring of estrous cycle stages and FACS, KEW: qRT-PCR, RNAseq analysis and differential gene expression analysis); Methodology: n.a.; Project administration: KEW, NH and RvA coordinated the planning and execution of the study; Resources: JJ provided access to the NKI Genomics Core facility; Software: n.a.; Supervision: RvA; Validation: KEW, NH and RvA evaluated and interpreted the data; Visualization: NH designed and prepared most figures; Writing - original draft: NH wrote the original draft as part of her PhD thesis with input from KEW and RvA; Writing - review and editing: RvA prepared the manuscript for submission and publication with input from all authors.

---

**Supplementary Fig. 1.**
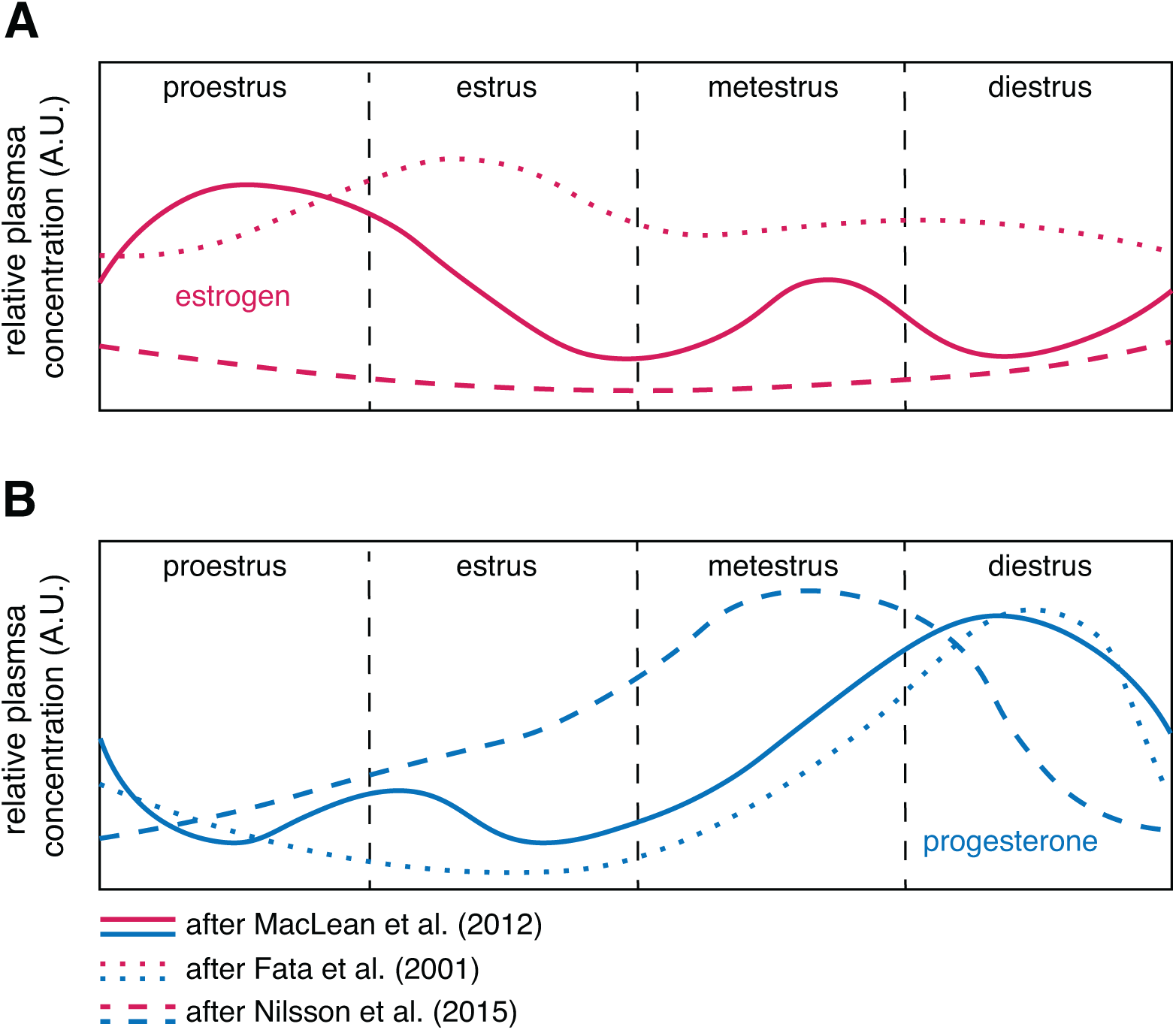
Estrogen and progesterone fluctuations during the mouse estrous cycle. Cartoon depicting relative hormone plasma concentrations during the four stages of the mouse estrous cycle as reported in three different studies [13–15]. Source data available via https://osf.io/xv83g/ (SuppFig1_sourcedata.xlsx). A) The highest estrogen levels occur in proestrus/estrus. B) The highest progesterone levels occur in metestrus/diestrus. A.U. = arbitrary units.

**Supplementary Fig. 2.**
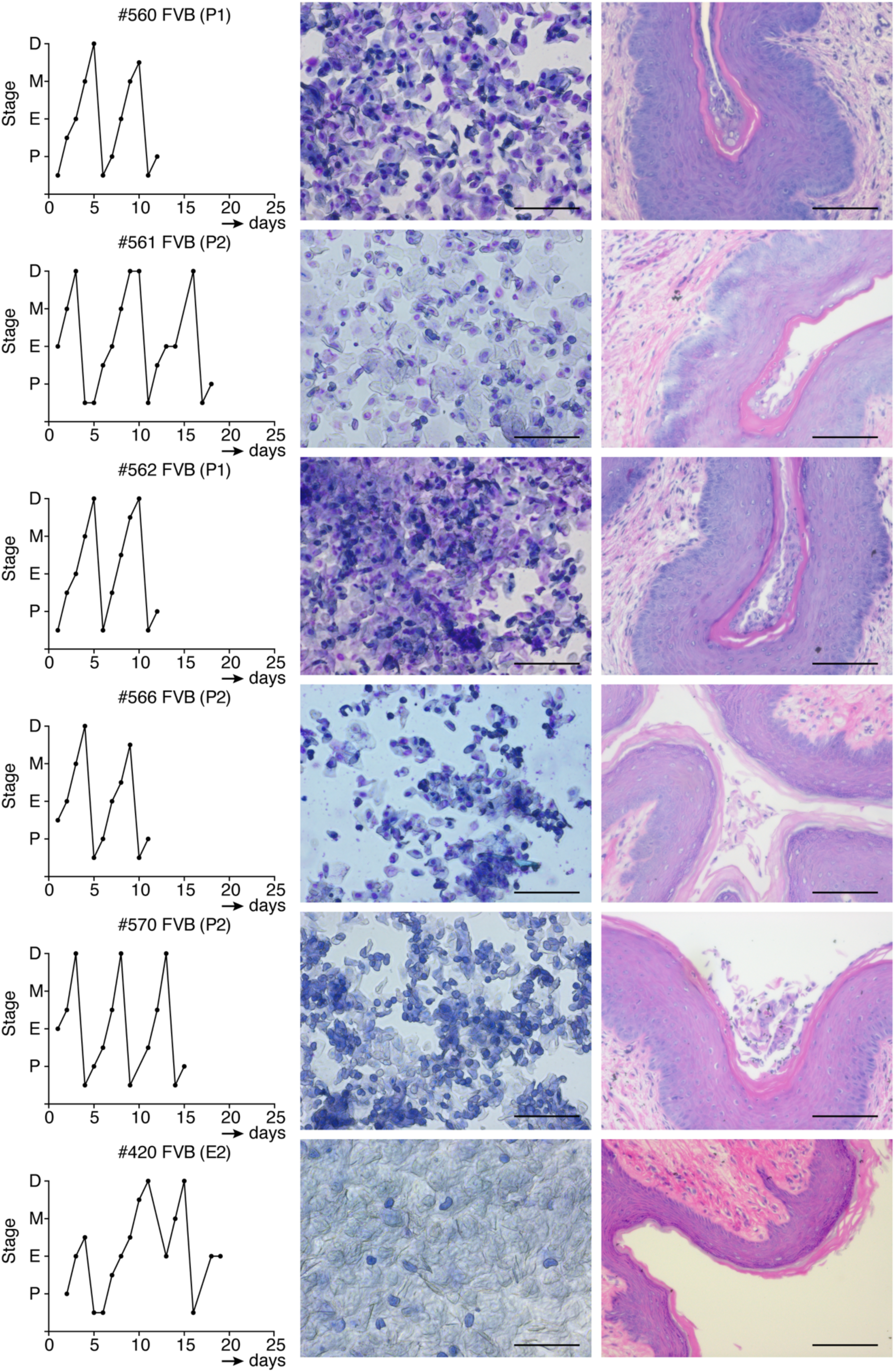
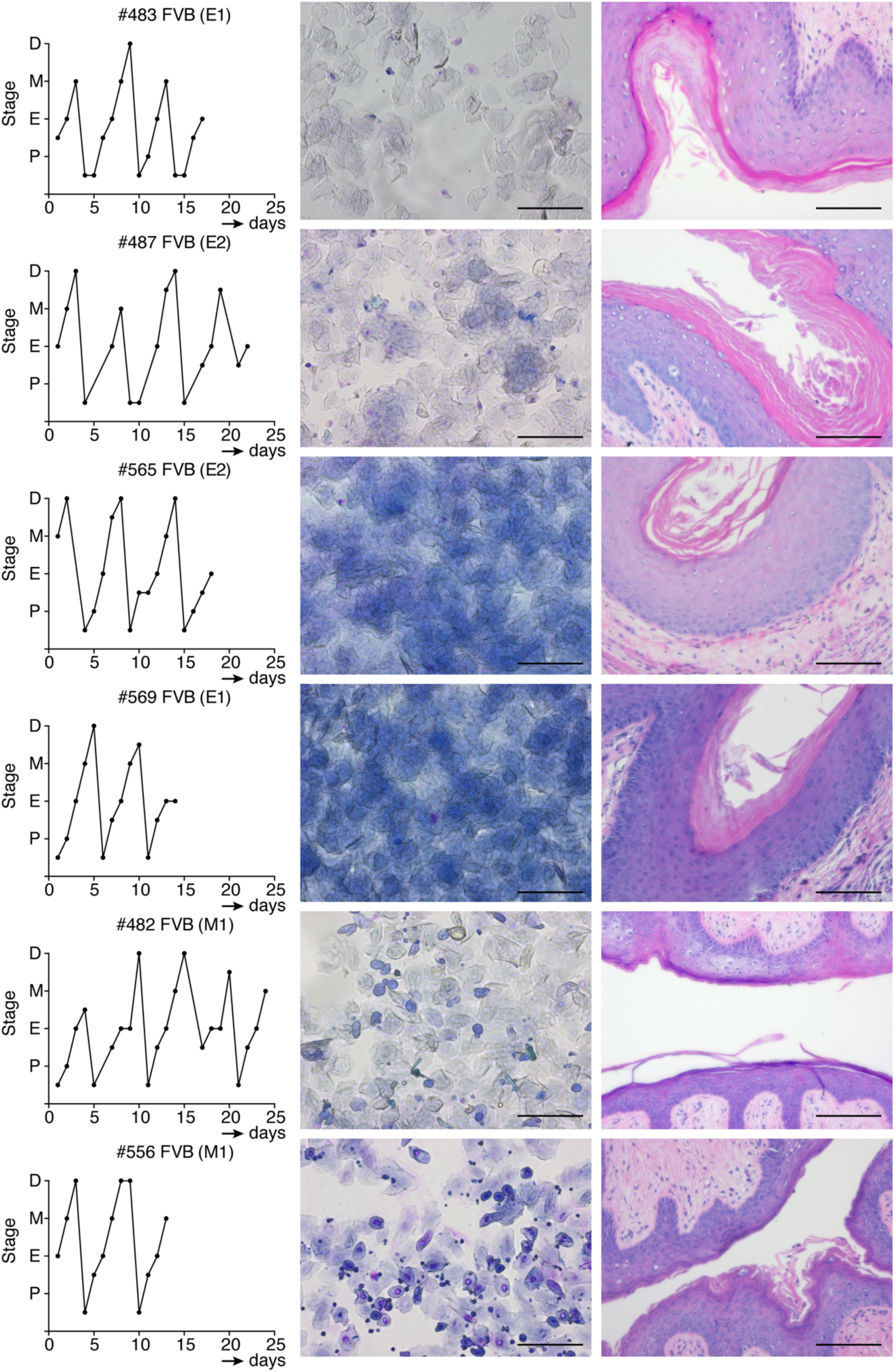
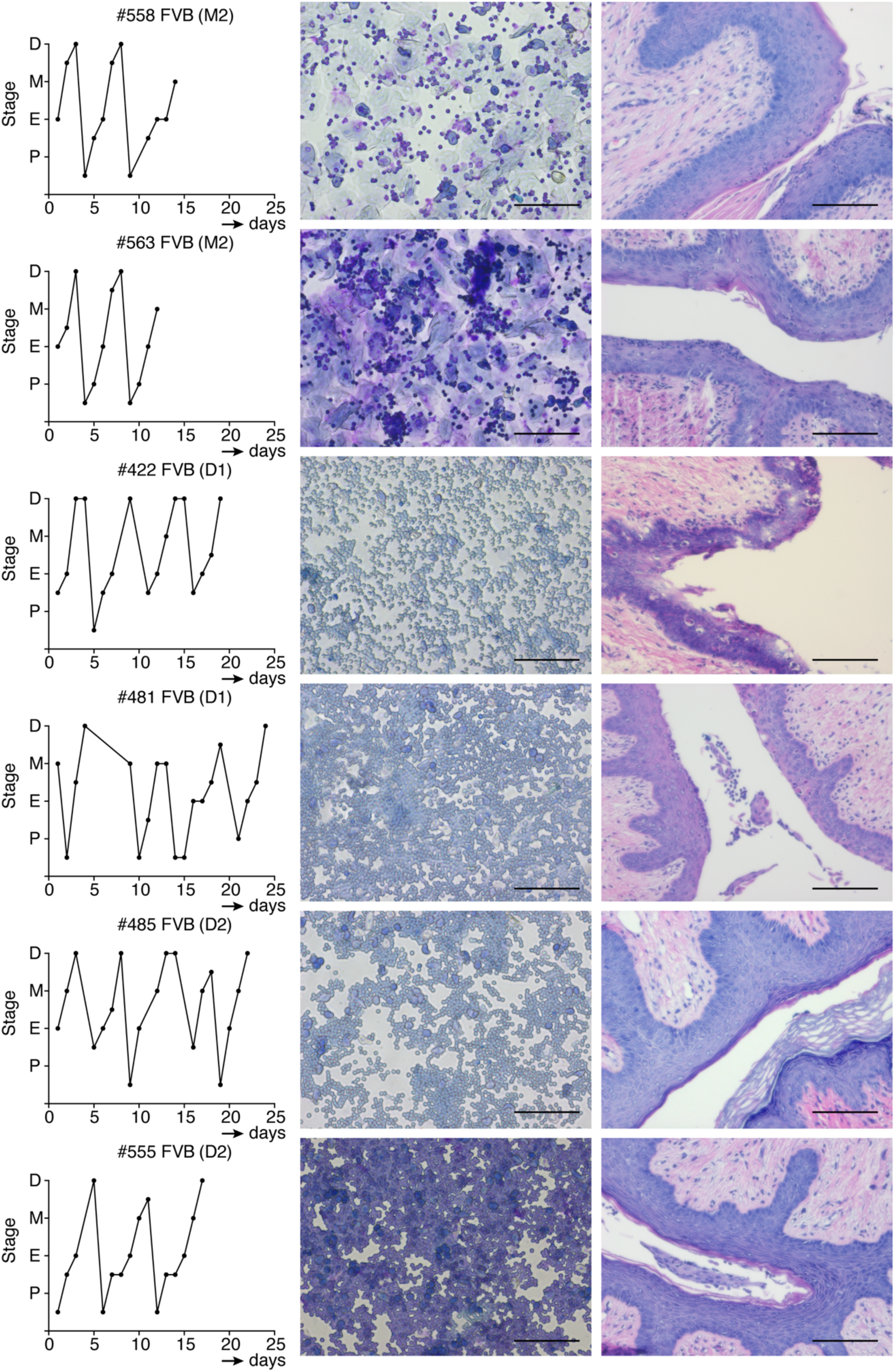
Estrous cycle monitoring of adult FVB/N mice. All mice that showed a stable estrous cycle and were selected for RNA-sequencing are shown in this figure, including the mice depicted in Fig.1. The left column shows estrous cycle monitoring over several days. The middle column and right column depict vaginal cytology and histology samples respectively, from the day of mammary gland isolation. Scale bar = 100 µm.

**Supplementary Fig. 3.**
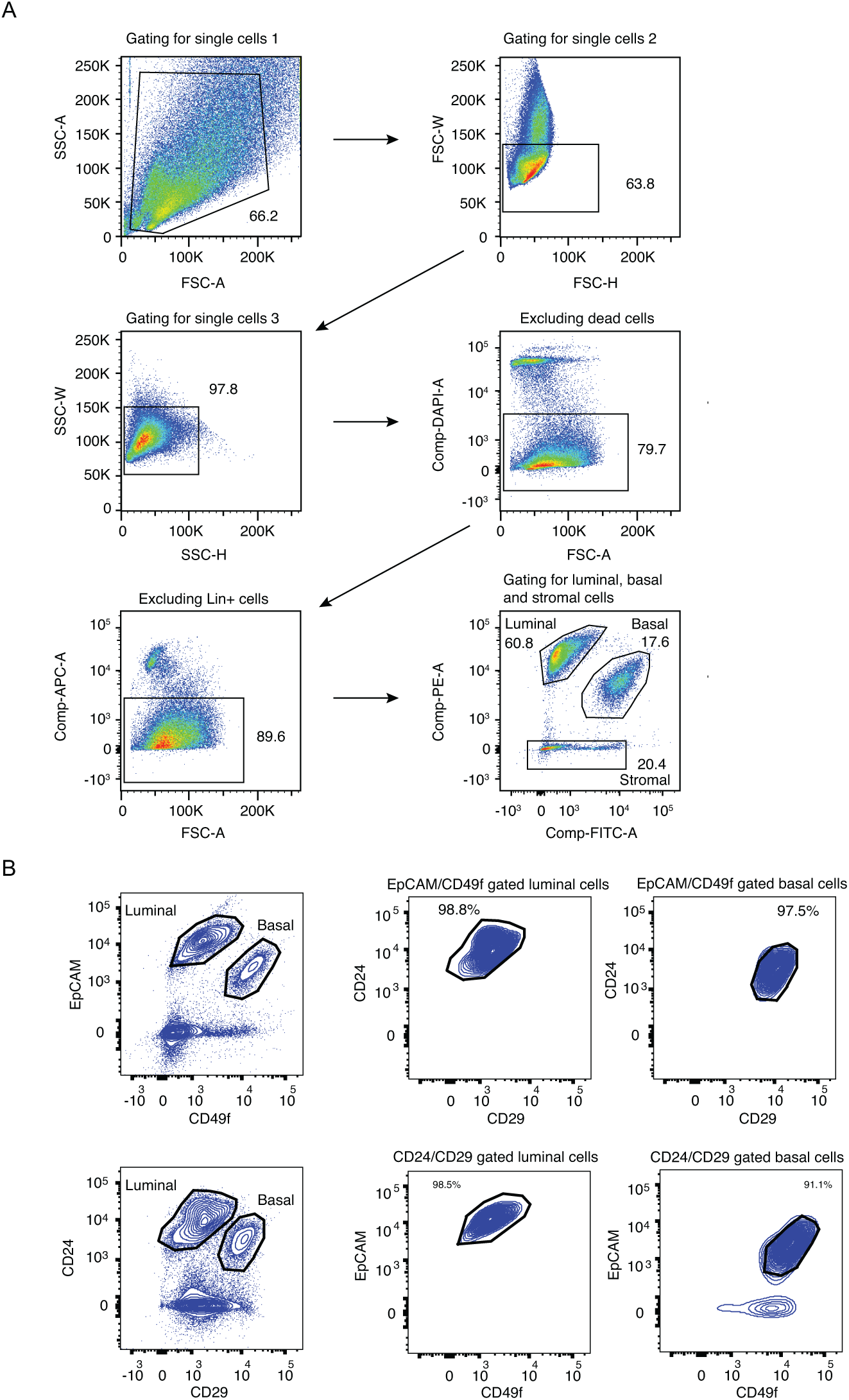
FACS strategy. A) Gating strategy for sorting luminal, basal and stromal mammary cell populations. SSC-A/FSC-A, FSC-W/FSC-H and SSC-W/SSC-H plots show gating for living single cells and to exclude dead cells, debris and doublets. DAPI plotted against FSC-A was also used to exclude dead cells. In the APC/FSC-A plot we gate for Lin-cells that are negative for CD45, CD31 and Ter119. CD45 and Ter119 markers were used to exclude haematopoietic cells and CD31 to exclude endothelial cells. EpCAM-PE and CD49f-FITC staining allowed us to gate for mammary luminal, basal and stromal cells before sorting. B) Direct comparison of the CD24/CD29 and EpCAM/CD49f staining. In our hands, EpCAM/CD49f gives slightly better separation of basal and luminal populations, making it our staining method of choice.

**Supplementary Fig. 4.**
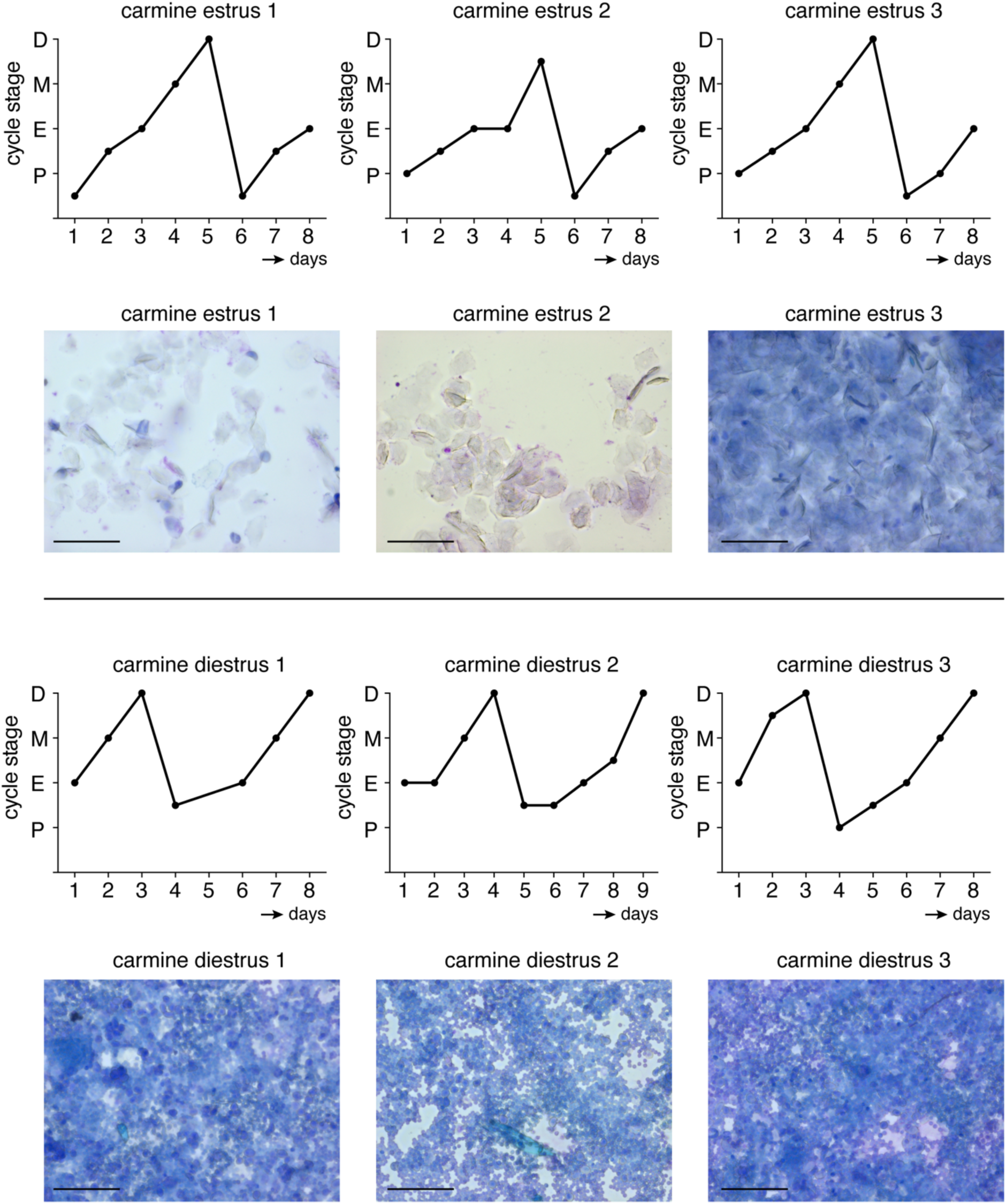
Estrous cycle monitoring of adult FVB/N mice for whole mount carmine staining. N=6 stably cycling mice were selected for carmine staining, 3 in estrus and 3 in diestrus. This figure shows the graphs of monitoring the estrous cycle for at least one week using vaginal cytology. The shown image is the cytology sample on the day of mammary gland isolation. Scale bar = 100 µm. Samples 1, 2, and 3 correspond to the same numbers in Supplementary Figure 5.

**Supplementary Fig. 5.**
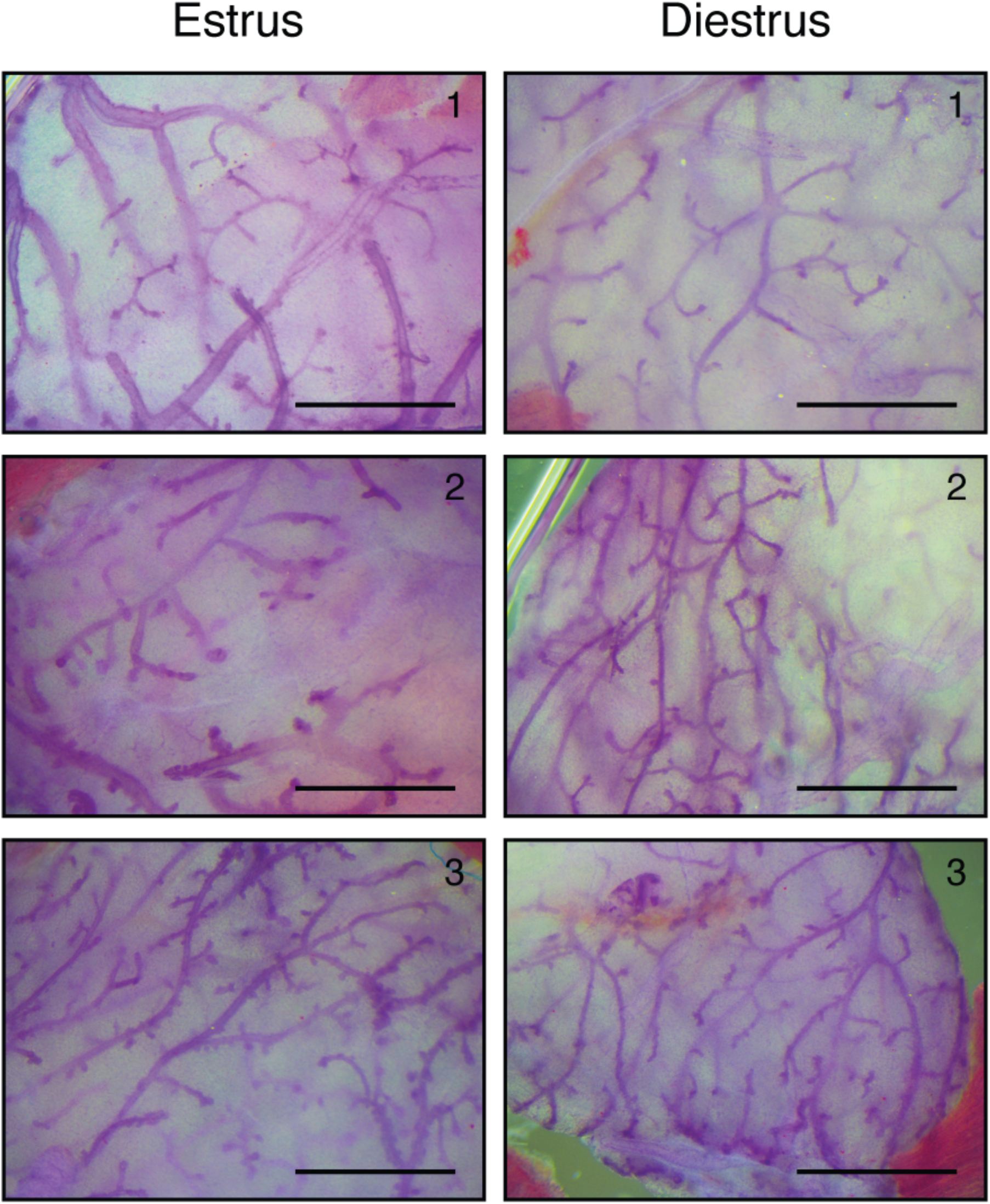
Whole mammary glands in estrus and diestrus show similar epithelial morphology. Images of carmine stained wholemount #3 mammary glands from n=6 different, estrous cycle staged mice (n=3 in estrus and n=3 in diestrus). Scale bar = 1 mm. Corresponding graphs of the estrous cycle monitoring of these 6 mice are provided in supplementary fig. 4.

**Supplementary Fig. 6.**
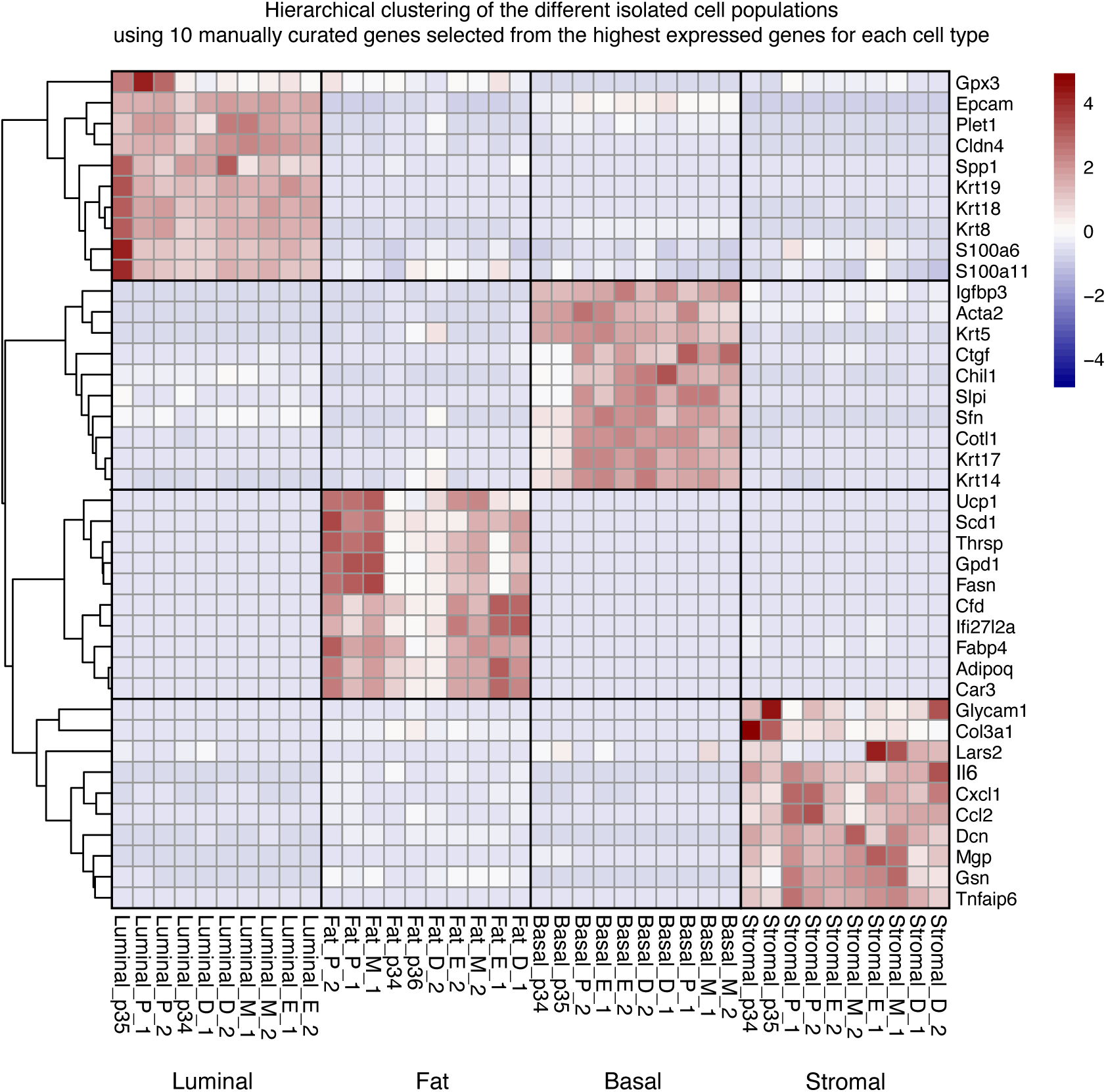
Unsupervised hierarchical clustering confirms proper isolation of the different cell populations. Heatmap showing expression (RPKM values, z-scale) of 40 genes (10 characteristic markers of each cell population) that were manually selected from the highest expressed genes across the different puberty and adult cell populations (top 35 for luminal, top 100 for basal, top 45 for fat and top 25 for stromal). P, E, M and D: estrous cycle stages. 1 and 2: adult RNAseq replicates. p34, p 35 and p36: puberty RNAseq replicates according to the samples in Table 1.

**Supplementary Fig. 7.**
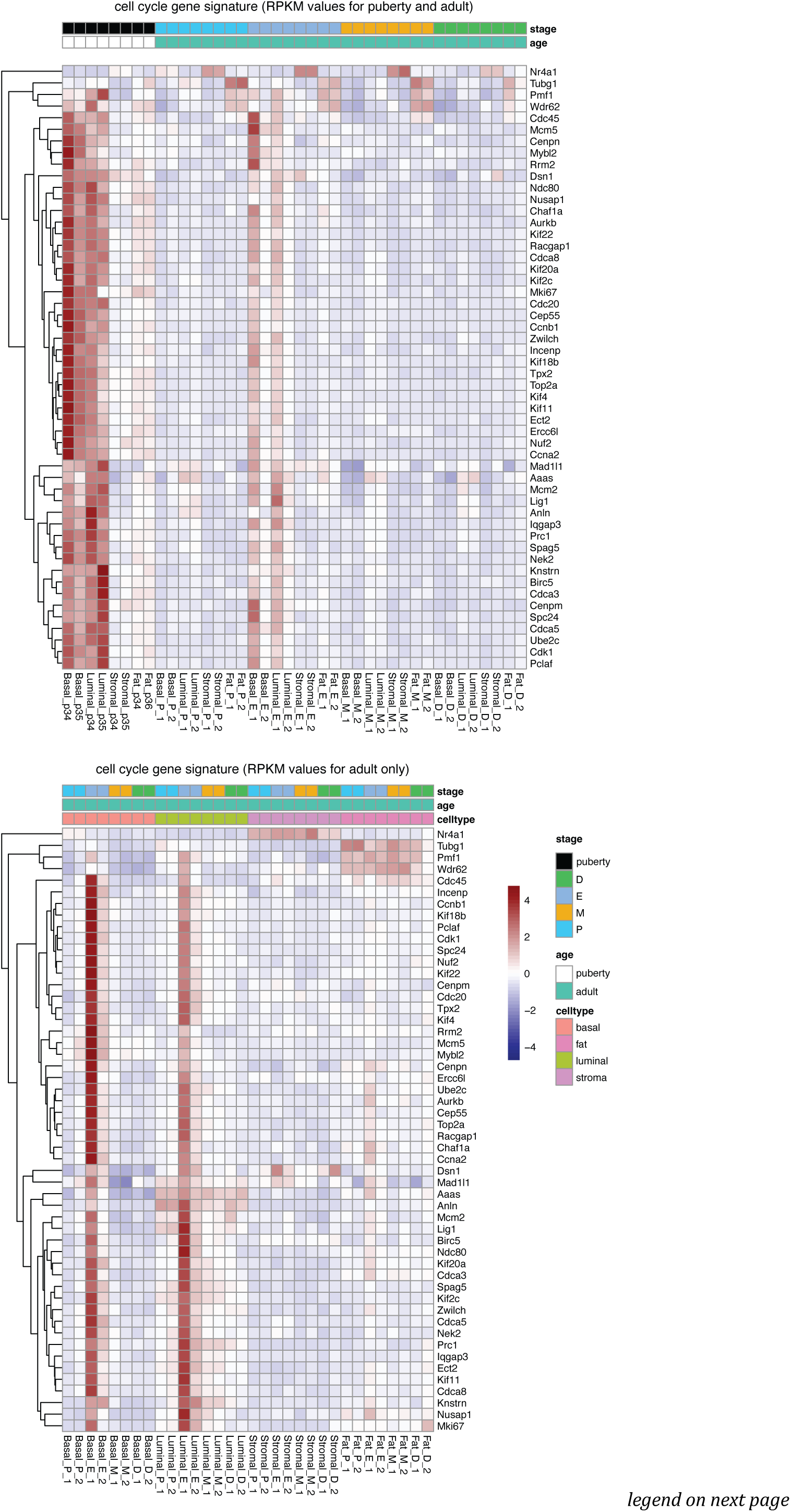
Estrous cycle dependent changes in cell proliferation genes. Heatmap showing expression (RPKM values, z-scale) of cell division genes in epithelial, but not in non-epithelial cell populations. A total of 85 genes showed differential expression based on pairwise comparisons of different estrous cycle stages in basal cells (FDR<0.01, Fig. 5B). Of these 85 genes, 65 matched with the GO biological process *cell cycle* (https://www.gsea-msigdb.org/gsea/msigdb/mouse/geneset/GOBP_CELL_CYCLE.html). Expression of these 65 genes was plotted for each RNAseq replicate (1, 2) across the pubertal and adult populations (**a**) or across the different adult cell populations (**b**).

**Supplementary Table 1.**
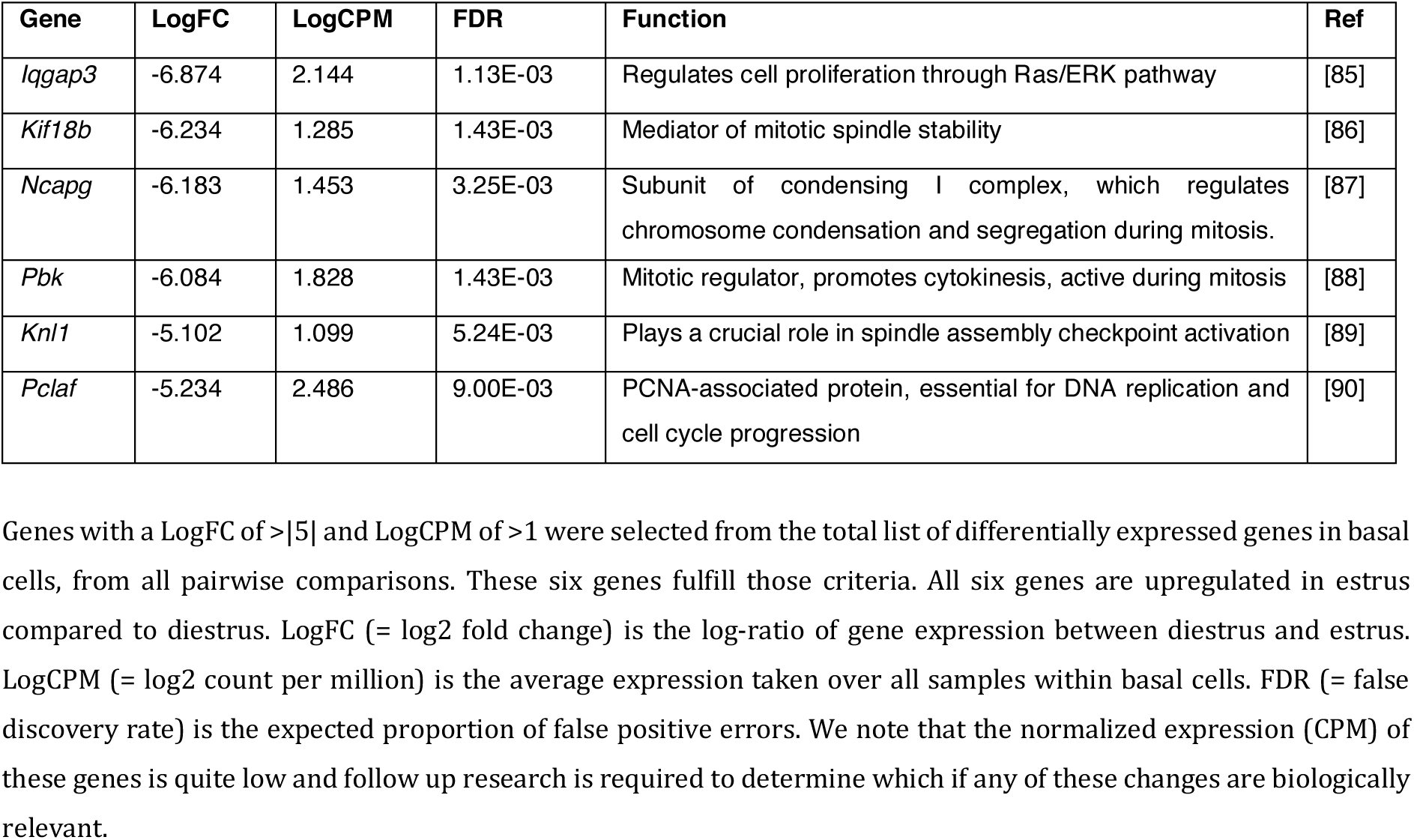
Differentially expressed genes with highest logFC in basal cells are directly linked to the cell cycle.

**Supplementary Table 2.** Differentially expressed Wnt genes during estrous cycle.

| Cell type | Comparison | Wnt gene | logFC | logCPM | FDR |
| --- | --- | --- | --- | --- | --- |
| Adipocytes | D vs P | *Wnt7b* | 3.724 | 5.529 | 4.90E-04 |
|  |  | *Wnt5b* | 2.362 | 6.136 | 9.42E-04 |
|  |  | *Wnt4* | 2.016 | 4.145 | 8.10E-03 |
|  |  | *Wnt10a* | 3.490 | 3.167 | 1.20E-02 |
|  | D vs E | *Wnt5b* | 2.268 | 6.136 | 1.39E-03 |
|  |  | *Wnt7b* | 2.800 | 5.529 | 3.55E-03 |
|  | D vs M | *Wnt7b* | 3.475 | 5.529 | 1.28E-03 |
|  |  | *Wnt4* | 2.395 | 4.145 | 4.10E-03 |
|  |  | *Wnt10a* | 3.486 | 3.167 | 1.78E-02 |
|  |  | *Wnt5b* | 1.458 | 6.136 | 2.60E-02 |
| Stromal | D vs E | *Wnt6* | -1.630 | 1.450 | 4.42E-02 |
|  | D vs M | *Wnt5a* | -1.504 | 5.603 | 4.11E-02 |

Only in comparisons of adipocytes and stromal cells differentially expressed (FDR <0.05) *Wnt* genes were present. Column "Comparison" shows the different pairwise stage comparisons that showed differentially expressed *Wnt* genes. P = proestrus, E = estrus, M = metestrus, D = diestrus. LogFC (= log2 fold change) is the log-ratio of *Wnt* expression between two different stages. LogCPM (= log2 count per million) is the average expression taken over all samples within a cell type. FDR (= false discovery rate) is the expected proportion of false positive errors. We note that the normalized expression (CPM) of these genes is quite low and follow up research is required to determine which if any of these changes are biologically relevant.

